# *De Novo* Design of Flavoproteins with Controlled Excited State Dynamics

**DOI:** 10.64898/2026.10.08.757584

**Authors:** Ethan L. Bungay, Juno Underhill, Meiyue Liu, Alexandra N. Stuart, Catherine R. Back, Jessica C. Berrones-Reyes, Paulina Dubiel, James A. Birrell, Aditya G. Rao, Christopher Williams, Rob Barringer, Philip Hinchliffe, Bruce R. Lichtenstein, Basile F. E. Curchod, Julea N. Butt, Thomas A. A. Oliver, Adrian J. Mulholland, J. L. Ross Anderson

## Abstract

Designing photochemical properties in proteins is a frontier challenge in biomolecular design, which, when realised, will open new avenues in catalysis and biomolecular electronics. Here we demonstrate an integrated computational design framework, supported by ultrafast spectroscopy, combining protein design, quantum mechanical/molecular mechanics molecular dynamics (QM/MM MD) and electronic structure calculations, to control flavin excited states in *de novo* proteins. Iterative design yields FC-4D2, a flavocytochrome with covalently bound flavin and non-covalently bound heme B and Fl-4D2, where the heme-binding site is removed. Both proteins are highly thermostable, and access all physiologically relevant flavin redox and protonation states, while high-resolution X-ray crystallography confirms the near-Ångström-level accuracy in design. Incorporating QM/MM MD and linear-response time-dependent density functional theory (LR-TDDFT) into the design workflow enables engineering of a flavin–tryptophan charge-transfer variant, Fl-4D2_CT_, that modifies the flavin absorption spectrum, substantially alters its redox properties and suppresses fluorescence. Ultrafast studies reveal how protein design reshapes the excited state relaxation timescale and mechanism. Our study establishes a modular strategy for programming flavin photochemistry in designed proteins.

## Introduction

Nature’s photoactive proteins provide an exquisite illustration of how the protein environment can precisely shape and modulate the photochemical and biophysical properties of a bound chromophore, tuning electronic absorption spectra, excited state lifetimes, and redox reactivity to ultimately suit the phenotypic demands of the organism.^1–5^ Of natural photoactive chromophores, flavins are perhaps the most versatile, participating in one-electron transfer, stepwise or concerted two-electron transfer, electron bifurcation, and direct photocatalysis.^6–8^ Flavins thus serve as the active, function-imparting cofactors in cryptochromes, LOV (light oxygen voltage) and BLUF (blue light using flavin) domains, and photolyases.^6,9^ More generally, flavins also play essential catalytic roles in many enzymes and proteins that do not functionally rely on light, performing diverse chemistries in monooxygenases, dehydrogenases, oxidases, dehalogenases, halogenases and reductases, while facilitating electron shuttling in electron transfer proteins such as flavodoxins.^10–13^

Significant advances have been gained through studying natural flavoproteins, however, fully accessing and exploiting this exceptional versatility in custom-built *de novo* designed proteins remains an outstanding challenge. As with other redox cofactors, many earlier attempts to create wholly *de novo* flavin-binding proteins have suffered from structural heterogeneity and poor thermostability, preventing exploration of the full photochemical functional landscape.^14,15^ Therefore, a strategy for *de novo* flavoprotein design with atomic accuracy, and the capability to program specific photophysical outcomes from first principles, has not yet been demonstrated.

The modular *de novo* redox protein platform developed by Anderson and coworkers offers an ideal platform for addressing this challenge.^16^ The four-helix bundle protein, 4D2, originally designed to recapitulate diheme electron transfer motifs from transmembrane respiratory complexes within a *de novo* scaffold, is expandable, modular and biophysically tuneable: heme redox potentials can be shifted predictably by mutagenesis or porphyrin replacement; cofactor-binding modules can be inserted or removed with minimal redesign; and the protein is amenable to surface-redesign and insertion into the inner membrane of *E. coli*.^16–19^

Here we develop and demonstrate an integrated computational–experimental pipeline for the design and mechanistic characterisation of photoreactive *de novo* flavoproteins, with full control over photochemical properties. Iterative deep-learning-guided sequence design using LigandMPNN combined with ESMFold structure prediction, yielded FC-4D2 and Fl-4D2, a *de novo* flavocytochrome and flavoprotein respectively, with both structures validated by X-ray crystallography. We extended the pipeline to incorporate QM/MM molecular dynamics and linear response time-dependent density functional theory (LR-TDDFT) spectral calculations, enabling the rational design of Fl-4D2_CT_, a flavin–tryptophan charge-transfer complex with markedly transformed photochemical dynamics compared to the parent scaffold. Ultrafast spectroscopy reveals the mechanistic origins for the dramatically different fates of the photoexcited states. Combined, we have established a generalisable programmable framework for controlling flavin photochemistry within structurally resolved *de novo* proteins. This represents a pivotal step in unlocking sophisticated light-activated bioenergetic circuitry in fully *de novo* designed proteins.

## Results and discussion

### Machine-learning-guided design of de novo flavoproteins

We developed a thermostable redox-active *de novo* flavoprotein starting from the previously reported monoheme four helix bundle m2-4D2^18^, with an internal cysteine residue for covalent flavin attachment as inspired by Bialas *et al*^15^, Fig.1a. Our initial approach involving Rosetta-based sequence redesign yielded structures that, despite high expression and solubility, displayed structural disorder at the flavin site, as exemplified by NMR spectra, Fig.S1. To address these issues, we subsequently constructed an extended 140-residue homologue of m2-4D2 comprised of three modules: the corresponding heme B binding module, a central cofactor-free module to enhance thermal stability, and a flavin module containing a core-facing cysteine for covalently appending 8-demethylriboflavin (8-dMeRF) via a thioether bond. This provided a basis for iterative ML-assisted redesign.

This extended starting scaffold (pTM = 0.843; ESMFold) was subjected to five rounds of iterative sequence redesign (Fig.1b), using LigandMPNN^20^ to design the flavin binding site and ESMFold^21^ to assess the apoprotein structures. In each round, the highest pTM structure was fed into the subsequent LigandMPNN redesign stage, with successive rounds of evolution increasing the mean pTM from 0.822 to 0.897 and mean pLDDT from 85.4 to 91.7, converging to a highly homologous population by iteration 5 with 64% sequence identity to the input, indicative of substantial sequence diversification (Fig.1c,d). The top-scoring sequence, designated FC-4D2 (flavocytochrome-4D2; pTM = 0.899, pLDDT = 91.9), and a heme-free version incorporating 9 mutations previously used to repack a heme-binding site^16^, Fl-4D2 (pTM = 0.895), were assessed by MM MD simulations prior to experimental characterisation. FC-4D2 and Fl-4D2 showed backbone RMSD values of 1.2 ± 0.2 Å and 1.4 ± 0.2 Å respectively over 5 × 50 ns trajectories, confirming structural stability, Fig.S5a,d. The isoalloxazine ring of the flavin remained anchored throughout these simulations through sustained hydrogen bonds from Arg37, Glu38, and Arg41 to the O4 locus and the ribityl tail, Fig.S5f. The heme-to-flavin edge-to-edge distance in FC-4D2 was 19.3 ± 0.6 Å (Fig.S5c), and theoretically capable of slow (millisecond) inter-cofactor electron transfer (ET).^22^

### Structural validation at atomic resolution

We recombinantly expressed FC-4D2 and Fl-4D2 in *Escherichia coli* from a standard pET-21(+) expression vector and purified both proteins *via* nickel affinity and size exclusion chromatography. To assemble the holoproteins, we subsequently loaded both proteins with cofactors *in vitro* and observed monodisperse peaks in the respective size exclusion chromatograms, Fig.S7,S8. Native ESI-MS demonstrated >95% flavination efficiency for both constructs. Circular dichroism spectroscopy confirmed the coiled-coil signature^23^ (MRE_222nm_/MRE_208nm_ ≈ 1) and hyperthermostability, with no sigmoidal unfolding transition observed up to 95°C for either design, Fig.2c-g. In addition, there was a notable divergence in the emissive properties of these designs; Fl-4D2 exhibits a prominent fluorescence emission spectrum upon 460 nm excitation, while FC-4D2 emits only weakly, due to a combination of heme absorption (a secondary inner-filter effect, Fig.S6) and an undetermined non-radiative decay pathway.

Protein crystals of both designs were successfully grown from sitting-drop vapour diffusion experiments. X-ray diffraction data were collected, and the structures determined using molecular replacement to resolutions of 1.35 Å (FC-4D2) and 1.8 Å (Fl-4D2) (Fig. 2i,j). The FC-4D2 and Fl-4D2 experimental structures align to the post-MM MD computational models with Cα RMSD of 0.844 Å and 0.840 Å, respectively, thus confirming the atomistic accuracy of the design methodology. The flavin attachment is characterised by a 1.8 Å thioether bond with a Cβ–S–C8 bond angle of 105°, with the attachment retaining the cofactor to high structural homogeneity, as exemplified in FC-4D2 in which electron density holes are visible in the isoalloxazine ring network. The FC-4D2 flavin module is more solvent accessible than the Fl-4D2 counterpart, likely due to a loss of helicity in the Ala30-Ala32 helix, a potential artefact of crystal packing (Fig.S9). In Fl-4D2, whereby this structure is retained, the flavin is within hydrogen bond distance to Ser31 at the N3 position, and within electrostatic interaction range to Arg37 at the N1-C2=O2 locus, as predicted through simulation. In FC-4D2, the heme– flavin distance of 19.8 Å further agrees with *in silico* predictions, whilst the heme module aligns with the equivalent site in the 4D2 experimental structure with 1.19 Å Cα RMSD, retaining the keystone secondary-coordination-sphere hydrogen bonds between histidines and partnering threonines, Fig.S10. Overall, these structures demonstrate the atomically precise design afforded by the iterative redesign approach and the amenability of these four-helix bundle modules to further redesign and rearrangement.

**Figure 1.**
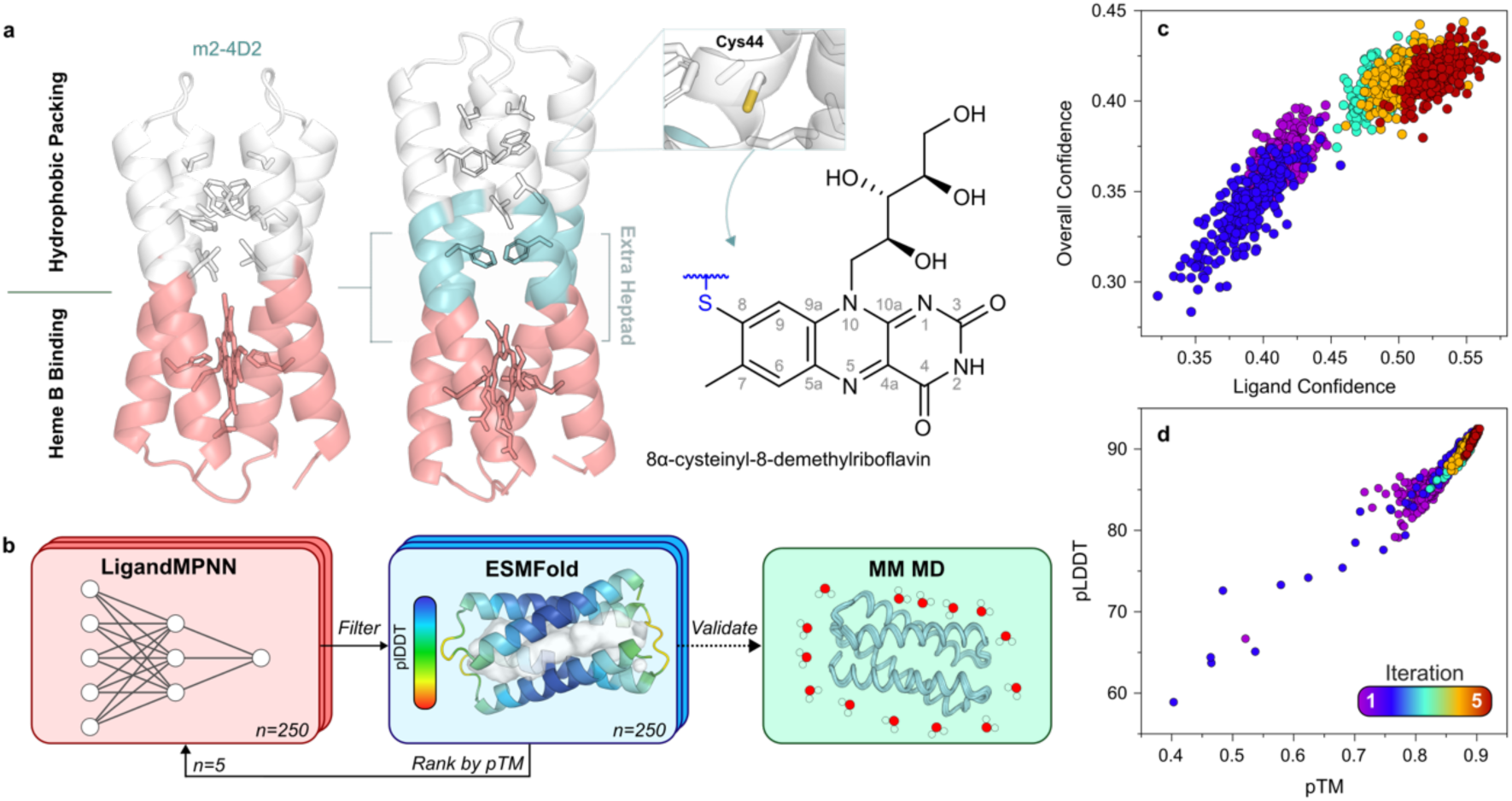
Iterative design towards a thermostable *de novo* flavocytochrome. (a) A three-module starting structure was rationally assembled from m2-4D2 (left) and additional four-helix bundle heptads, yielding a 140-residue maquette with an internal cysteine capable of the thioether linkage to 8-demethylriboflavin. (b) Iterative *in silico* directed evolution redesign pipeline, wherein the starting structure (a) was fed into 5 cycles of LigandMPNN-mediated sequence redesign and ESMFold apo structure prediction, before validation with MM MD. Ranking metrics used included LigandMPNN confidence (c) and ESMFold pTM and pLDDT (d).

**Figure 2.**
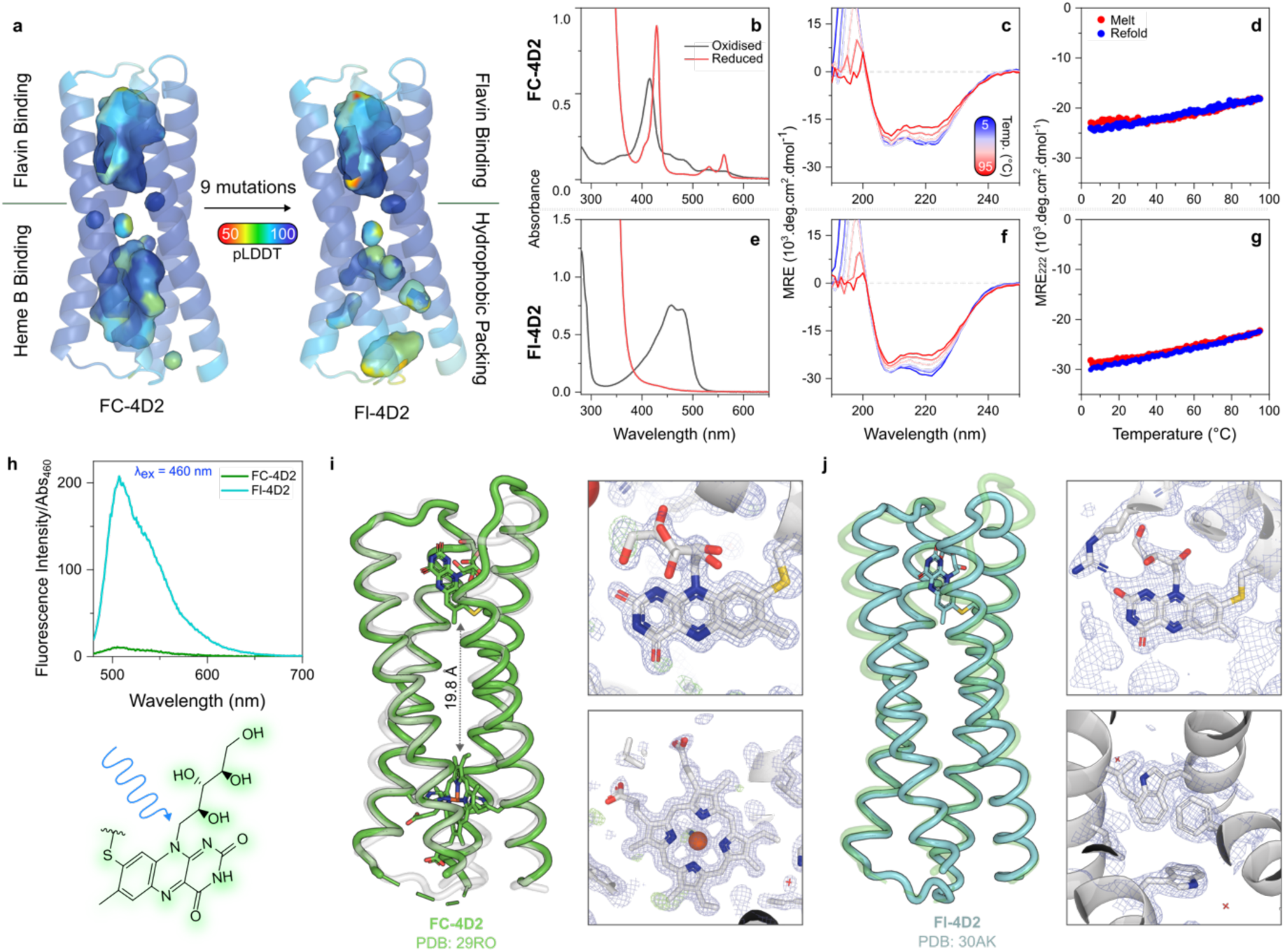
Biophysical characterisation of FC-4D2 and Fl-4D2. (a) ESMFold structure predictions of apo FC-4D2 and Fl-4D2, coloured by pLDDT with respective modules and ligand-binding cavities presented. (b) Steady-state absorption spectra of FC-4D2 in the oxidised (black) and reduced (red) states. (c) Far-UV CD spectra of FC-4D2 from 5-95 °C. (d) Temperature dependence of FC-4D2 CD signal at 222 nm during melt (red) and refolding (blue). (e-g) same as (b-d) for Fl-4D2. (h) Absorbance-corrected fluorescence emission spectra of FC-4D2 (green) and Fl-4D2 (cyan) upon 460 nm excitation. (i) Crystal structure of FC-4D2 (PDB: 29RO) resolved at 1.35 Å. (left) Tube representation of structure (green) overlaid with the post-MM MD designed state (white, semi-transparent). (right) 2F_o_-F_c_ weighted electron density (blue mesh, 1σ, 0.294 e^−^/Å^3^) and F_o_0–F_c_ weighted difference maps (green mesh, 3σ, 0.280 e^−^/Å^3^) overlaid on top of FC-4D2 cofactors. (j) Crystal structure of Fl-4D2 (PDB: 30AK) resolved to 1.8 Å. (left) Full structure of Fl-4D2 (cyan) aligned with FC-4D2 (green, semi-transparent), (right) electron density representations of the flavin and hydrophobic-packing modules, same conditions as (i).

### Electrochemical characterisation and pH-dependent redox tuning

To electrochemically characterise the bound redox-active cofactors in these designs, we first employed protein film electrochemistry (PFE), in which our proteins were adsorbed onto mesoporous indium tin oxide electrodes and subjected to cyclic voltammetry. Baseline-subtracted ‘Faradaic’ currents of Fl-4D2, representative of interfacial electron transport between the redox centre and electrode, were modelled as a sequential 2 × 1-electron redox process whereby *E_1_* and *E_2_* represent the midpoint potentials of the oxidised/semiquinone and semiquinone/hydroquinone transitions, respectively. The *E_1_*/*E_2_* potentials for the Fl-4D2 flavin, averaged across reducing/oxidising potentiometric sweeps, were −168.2 ± 7.5 mV and −196.3 ± 0.8 mV, squarely within the electrochemical range exhibited by natural flavoproteins (-400 to +60 mV vs SHE)^24^, Fig. 3b. With discrete potentials and an average half-height peak width (W_1/2_) of 83.7 ± 6.3 mV, this behaviour is more consistent with sequential one-electron transitions, *via* the intermediary semiquinone state, rather than concerted two-electron chemistry. We further validated that these potentials were representative of the protein in solution by performing optically transparent thin-layer electrochemistry (OTTLE),^25^ with equilibration for 30 min at each 20 mV applied potential. The first derivative of the resulting sigmoidal Nernst titration showed close agreement with the PFE-derived Faradaic currents.

**Figure 3.**
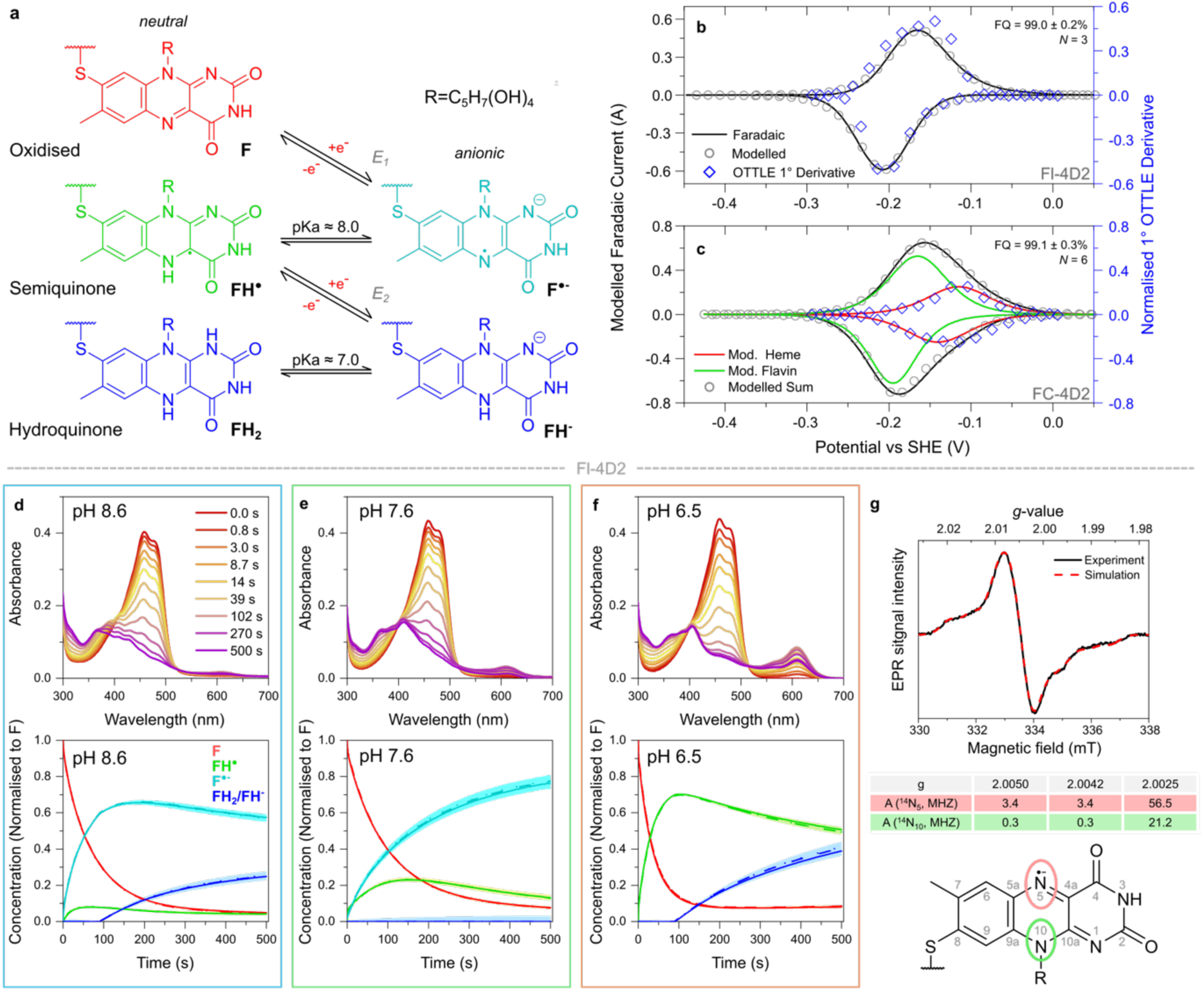
Electrochemical and photophysical analyses of FC-4D2 and Fl-4D2. (a) Five physiologically prevalent redox/protonation states of flavins. (b) Faradaic currents of Fl-4D2 from PFE at pH 8.6, 0.01 V.s^−1^ (black line), modelled with a single 2 × *n*=1 redox centre (grey circles) and overlaid with first derivative of OTTLE flavin titration (blue diamond). Fit quality (FQ) and number of repeats (N) presented on the plots. (c) Same as (b) for FC-4D2, but with Faradaic currents modelled with distinct *n=1* (‘Mod. Heme’, red) and 2 × *n*=1 (‘Mod. Flavin’, green) redox centres, with the sum presented as grey circles. Overlaid with first derivative OTTLE titration of the heme moiety (blue diamond) which aided delineation of Faradaic current contributions. (d-f) Continuous-wave (cw) photoreduction of Fl-4D2 at pHs 8.6–6.5. Steady-state absorbance spectra (top) following 0–500 s of illumination. Deconvolved spectral contributions from 4 flavin redox states (bottom). Shaded areas are a 95% confidence interval of the fit, and three repeat experiments are shown (full/dashed/dotted lines). (g) X-band (9.359 GHz) cw EPR spectrum of Fl-4D2 with a 100-fold excess of sodium dithionite freeze-quenched after 100 ms (black line), fitted with a simulated spectrum (red). (bottom) Structure of the isoalloxazine in anionic semiquinone state, with highlights representing the N5 and N10 loci likely responsible for the ^14^N hyperfine coupling, with parameters listed in the table.

**Figure 4.**
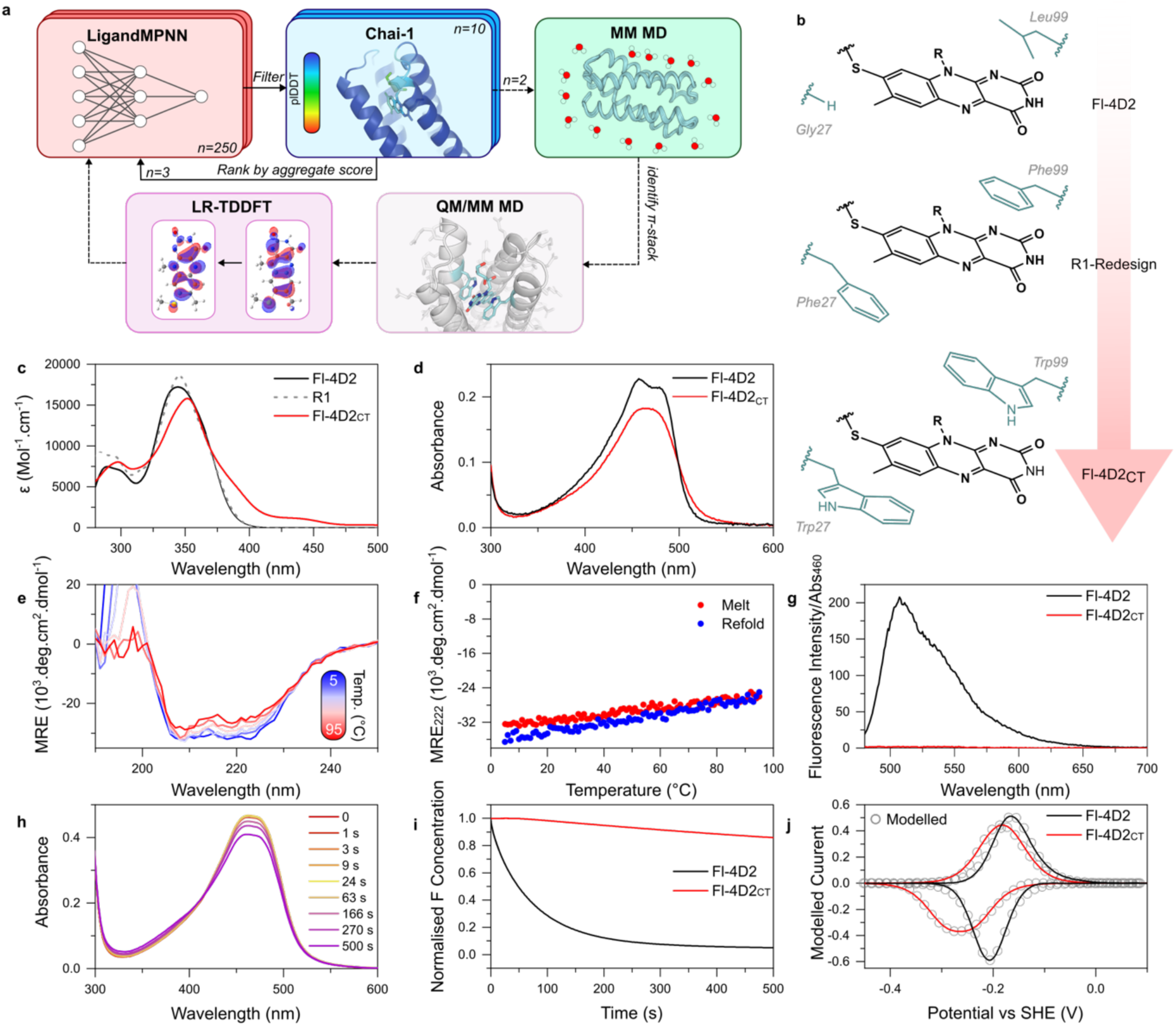
Design and characterisation of Fl-4D2_CT_. (a) QM/MM-integrated *in silico* directed evolution pipeline, modifying and extending the pipeline shown in Fig.1a to include ligand-protein co-diffusion with Chai-1, QM/MM MD of promising CT-like snapshots from MM MD, and LR-TDDFT to predict absorption spectra from the QM/MM MD trajectory. (b) Occupancy of flavin-sandwiching residue positions 27 and 99 in proteins of increasingly red-shifted predicted absorbance: Fl-4D2, R1-redesign, and Fl-4D2_CT_. (c) Predicted absorption spectra of Fl-4D2 (black), R1-redesign (grey dashed) and Fl-4D2_CT_ from LR-TDDFT calculations. (d) Steady-state absorption spectra of Fl-4D2 and Fl-4D2_CT_ at 5 μM 8-dMeRF obtained at pH 8.6 (as is each following plot). (e) Far-UV CD spectra of Fl-4D2_CT_ as a function of temperature, with a corresponding melt/refold curve recorded at 222 nm (f). (g) Absorbance-corrected fluorescence spectrum of Fl-4D2 and Fl-4D2_CT_ after 460 nm excitation, showing almost complete suppression of emission from Fl-4D2_CT_. (h) Steady-state absorbance spectra of Fl-4D2_CT_ following increasing cw irradiation. (i) Average normalised oxidised flavin (F) concentration of Fl-4D2 and Fl-4D2_CT_ across 500 s of cw white light illumination. (j) Modelled Faradaic PFE peaks of Fl-4D2 and Fl-4D2_CT_ obtained at 0.01 Vs^−1^.

For the multicofactor FC-4D2, the additive contributions of two redox centres with similar potentials resulted in a conjoined Faradaic peak, necessitating the use of the heme OTTLE derivative to identify the cofactor’s potential prior to subsequent modelling with equal contributions from an *n*=1 and a 2 × *n*=1 redox centre, Fig.3c. A fit quality, defined as 1 minus the percentage mean absolute deviation between experimental and modelled currents, above 99% supported the presence of both electroactive cofactors with the expected stoichiometry. The FC-4D2 heme is characterised by a single midpoint potential, corresponding to the Fe^3+^/Fe^2+^ transition, of −128.9 ± 3.7 mV, near-identical to that of m2-4D2.^18^ This suggests preservation of the local electrostatic environment of the heme module across design iterations. Furthermore, the FC-4D2 flavin exhibits a similar redox window to Fl-4D2, with *E_1_*/*E*_2_ values of −170.3 ± 1.6 mV and −200.4 ± 2.0 mV, respectively, and an average W_1/2_ of 85.2 ± 3.7 mV, further demonstrating that the scaffold can be modified without substantially perturbing the redox chemistry of adjacent modules.

### Accessing a range of flavin redox states

Following electrochemical identification of sequential single–electron chemistry in both designs, we examined the photoreduction kinetics of Fl-4D2 under continuous-wave (cw) white-light illumination, Fig.3f-h. After irradiation at pH 8.6, the flavin of Fl-4D2 is predominantly reduced to the anionic semiquinone (Fig.3a), with a small population of neutral semiquinone visible, as evident by the appearance of an absorption peak at 610 nm. The 2-electron reduced hydroquinone slowly accumulates from ∼90 seconds after the initial illumination, and after 500s 90% of the flavin has been transformed into either long-lived semiquinone or hydroquinone forms. The populations of these species and their lifetimes under cw illumination can be controlled with pH; at pH 6.5, the neutral semiquinone is the dominant one-electron reduced species, while at pH 8.6, the anionic semiquinone is favoured. From this measurement, the hydroquinone species is assigned as a mixture of the neutral and anionic species, as they cannot be readily distinguished by their visible absorption bands. With freeze-quenched EPR, we further explored the nature of the semiquinone species at pH 8.6. These measurements revealed a radical centred at *g* = 2.005, with a linewidth of 1 mT – consistent with the anionic form of the one-electron reduced flavin species^26^, and evidence of two ^14^N couplings (Fig.3g). Inclusion of ^1^H hyperfine coupling, either alone or as a second protonated species, yielded similar quality simulations. Thus, the EPR data supports the predominance of the anionic flavin semiquinone at pH 8.6, but cannot exclude the presence of the neutral form. Anionic semiquinone stabilisation could be attributed, as evident from the solved structure, to electrostatic interactions between Arg37 guanidinium and the isoalloxazine N1-C2=O2 locus, with charge-withdrawing interactions at this site often found in flavin-based electron bifurcating systems.^27,28^ Combined, this demonstrates we can control access to at least 4 of the 5 physiologically relevant flavin redox/protonation states in Fl-4D2, opening the door to downstream functional tuning and chemistry selection.

### QM/MM-guided design of a flavin–tryptophan charge-transfer (CT) complex

To engineer qualitatively distinct excited state dynamics in Fl-4D2, we pursued a charge-transfer (CT) complex with π-stacking between aromatic residues and the flavin isoalloxazine. This strategy aimed to enable fast electron donation from the aromatic side chain to the photoexcited flavin to produce a high yield of charge separation; with further design, this charge could be extracted and serve as a basis for *de novo* reaction centres.^4,29,30^ With our chosen approach, we anticipated our design would broaden the absorption spectrum, reduce absorbance cross-section, and suppress fluorescence, comparable to natural flavodoxins.^31–33^ Starting from the Fl-4D2 crystal structure, we targeted fourteen residues surrounding the flavin for redesign with LigandMPNN, biasing the primary coordination sphere towards aromatic amino acids. Of 250 generated sequences per cycle, the top ten candidates (based on overall/ligand confidence) were co-folded with covalently attached 8-demethylriboflavin using Chai-1^34^, and the highest confidence design (based on pTM/ipTM) served as the input back into LigandMPNN for 3 cycles of iterative design, Fig.4a. The highest confidence structure from these 3 cycles was then subjected to MM MD simulation to assess conformational stability and identify desirable π-stacking interactions between aromatics and the isoalloxazine ring of the flavin (Methods). QM/MM MD simulations, using DFT/wB97X-D3/def2-SVP within the AMBER/TeraChem^35–39^ interface, were initiated from the selected π-stacking snapshot to generate a conformational ensemble with more highly accurate 8-demethylriboflavin structures. 100 trajectory snapshots from QM/MM MD were then extracted for LR-TDDFT/TDA/wB97X-D3/def2-SVP calculations (for electronic structure benchmarking, see SI Section S5). The calculated excitation energies and transition dipole moments from LR-TDDFT were used to predict the absorption spectra, and to assess charge-transfer character to the nearest aromatic residues for each electronic transition. This methodology has proven successful in our recent combined experimental/theoretical study of the riboflavin chromophore in water.^40^

A first-round design (R1) with proximal phenylalanines produced no computed spectral modification for the main band (Fig.4c) and minimal CT character (Table S2, Fig.S17). The R1 snapshot with the largest CT character for the S_0_ – S_1_ transition was selected for another design round, generating 750 sequences, of which the top-scoring variant was designated Fl-4D2_CT_. This second design round introduced two tryptophans in place of the phenylalanines, sandwiching the isoalloxazine ring. Fl-4D2_CT_ showed greater conformational stability than R1 in the MM MD (Fig.S15), and LR-TDDFT calculations from the QM/MM MD simulation predicted a reduced molar extinction coefficient, and a longer-tailed absorption in the long wavelength region with significant CT character, Fig.4c, Table S2, Fig.S16.

### Experimental characterisation of CT character and redox properties

We expressed Fl-4D2_CT_ using the conditions and protocols established for Fl-4D2. Purification confirmed monodispersity with size exclusion chromatography, yet with a reduced efficiency of flavination (68%) compared to Fl-4D2 (>95%), Fig.S19. CD spectroscopy confirmed the retention of the expected secondary structure and hyperthermostability, with minimal loss of helicity up to 95°C, Fig.4e,f. As predicted, the steady-state absorption spectrum of Fl-4D2_CT_ differs markedly from Fl-4D2, displaying a reduced absorbance and some broadening, with the loss of vibronic structure evident for Fl-4D2 at 457 and 480 nm, Fig.4d. This is consistent with observations of flavodoxin proteins where charge-transfer in flavins is dominant.^33^ Fl-4D2_CT_ has no steady state fluorescence, and does not undergo significant photoreduction upon cw white-light irradiation, thus exhibiting enhanced photostability, Fig.4g-i. This contrasts Fl-4D2’s significant fluorescence yield and marked generation of semi/hydroquinone states upon photoreduction.

Deviations were also noted in the redox behaviour of the bound flavin. We modelled PFE-derived Faradaic currents as sequential 2 × 1-electron processes, revealing *E_1_*/*E_2_* potentials of -197.3 ± 5.6 and -253.1 ± 3.8 mV vs SHE, average respective shifts of -29 and -57 mV against Fl-4D2. The significant shift in *E_2_* potential is indicative of flavodoxin-like behaviour in Fl-4D2_CT_, whereby preferential tryptophan-stabilisation of the oxidised ground state decreases the midpoint potential of the semiquinone/hydroquinone pair.^41,42^ The increased peak separation is reflected in the W_1/2_, especially in the reductive direction (Fl-4D2_CT_ W_1/2_^Red^ = 129.7 ± 5.2 mV, Fl-4D2 81.0 ± 6.7 mV), further emphasising the kinetic bottleneck to hydroquinone formation. The increased peak separation of Fl-4D2_CT_ mimics but does not recapitulate stereotypical flavodoxin behaviour, with natural systems exhibiting peak separations of greater than 300 mV and *E_2_* potentials as low as –400 mV vs SHE.^43,44^ These are jointly achieved *via* neutral semiquinone stabilising interactions at the flavin’s N5 locus, alongside anionic hydroquinone-destabilising interactions through acidic residues in the flavin pocket, both of which were absent across 100 ns MM MD of Fl-4D2_CT_.^45^

### Elucidating flavoprotein excited state relaxation mechanisms

To rationalise the different fluorescent and photoreduction properties of Fl-4D2 and Fl-4D2_CT_ flavoproteins, we used ultrafast transient absorption (TA) spectroscopy. Both flavoproteins were excited with 480 nm laser pulses using an established laser system.^40^ Key differences between the two flavoproteins are apparent in TA data (Fig.5) at longer probe wavelengths (>500 nm), and on >10 ps timescales (full analysis is given in the SI, Section S3). At the very earliest time delays, Fl-4D2 shows stimulated emission at 550 nm from the S_1_ state consistent with steady-state fluorescence (Fig.4f). This feature is entirely absent in Fl-4D2_CT_, and instead, a broad photoinduced absorption (500–700 nm) is produced within the instrument response (146 fs) resembling the known tryptophan radical cation (W^•+^) absorption.^46–50^ Immediate appearance of the W^•+^ absorption indicates very fast electron transfer from the sandwiched tryptophan residues in Fl-4D2_CT_, or the direct formation of a flavin anion–tryptophan cation (F^•–^W^•+^) CT state delocalised over both moieties (SI Section S3 for discussion of spectral assignment), the latter predicted by QM/MM calculations. The F^•–^W^•+^ state is short-lived, and rapid relaxation back to the ground state is observed within 50 ps, similar to natural flavodoxins,^31,32^ thus preventing any sustained reduction.

**Figure 5.**
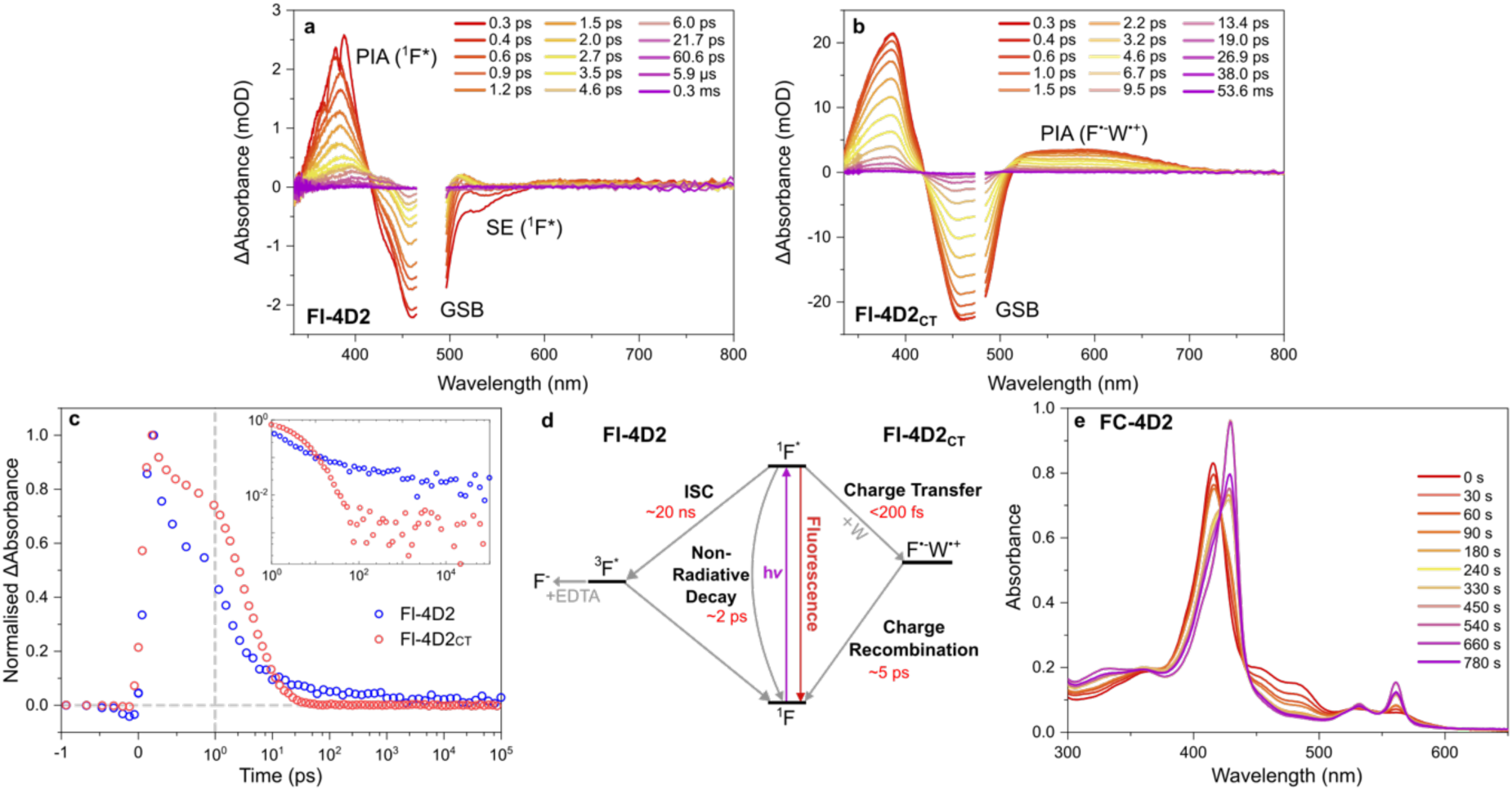
Ultrafast spectroscopy determines the excited state decay pathways in Fl-4D2 and Fl-4D2_CT_. Transient absorption spectra of (a) Fl-4D2 and (b) Fl-4D2_CT_ at selected time delays, highlighting photo-induced absorption (PIA), ground state bleach (GSB) and stimulated emission (SE) transient features. (c) Comparison of kinetics at 400 nm for Fl-4D2 and Fl-4D2_CT_. The vertical dashed line indicates a change from linear to logarithmic time axis, and the inset illustrates the data on a logarithmic signal intensity axis. (d) Schematic illustrating the different excited state relaxation pathways in the two *de novo* flavoproteins. (e) Steady-state absorbance spectra of FC-4D2 after irradiation with 480 nm pulsed laser light (10 kHz repetition rate).

In contrast, Fl-4D2 does not show complete ground state bleach (GSB) recovery even up to time delays of 400 µs (Fig.5c, Fig.S37-S39). Instead, a small portion of excited states in Fl-4D2 undergo intersystem crossing (ISC) to a long-lived triplet state (Fig.5d), which must serve as a precursor to forming the flavin semiquinones and hydroquinones observed in photoreduction experiments (Fig.3f-h). Notably, ∼90% of the Fl-4D2 excited states relax to the ground state within 10 ps (Fig.5c), indicating a significant non-radiative pathway that limits the available reservoir of excited flavin molecules capable of photoreduction. The long-lived population in Fl-4D2 likely arises from a sub-ensemble for which this quenching pathway is not efficient, with photoexcited flavin molecules that instead ISC to a long-lived triplet state.

It is important to reconcile the observed photoreduction of Fl-4D2 under cw irradiation with the complete absence of these products in TA measurements using pulsed lasers. The high-fluence cw white-light source has the potential to provide multiple photons into the flavin photocycle, *e.g.* to excite flavin semiquinones products, which may be responsible for the full photoreduction (Fig.3). Hence, we additionally performed photoreduction experiments with a 10 kHz pulsed 480 nm laser source, to minimise the potential for sequential multiphoton-induced photochemistry. Full photoreduction was still observed under pulsed excitation (SI Section S2.2), indicating that the lack of photoreduction observed in TA is not due to a different mechanism, but that only a very small portion of excited states are reduced in an individual photocycle. Based on the photoreduction rate, we expect ∼0.02% of excited flavin molecules to be reduced per pulse, which is below the TA noise floor (∼1%, SI Section S2.3). Remarkably, this demonstrates that the *de novo* flavoprotein is sufficiently robust to undergo thousands of photocycles allowing for sufficient accumulation of the flavin semiquinone to be observed in steady-state spectroscopy, proving a critical species to facilitate downstream chemistry. Photoreduction of FC-4D2 was also demonstrated under pulsed irradiation conditions (Fig. 5e), where >50% of heme B in FC-4D2 was reduced by electron transfer from flavin semiquinones or hydroquinones over 500 s. Ultimately, ultrafast studies reveal that the photophysical function of the *de novo* flavoproteins are controlled in Fl-4D2_CT_ by either rapid electron transfer followed by fast back electron transfer, or in Fl-4D2 by intersystem crossing to form long-lived states that can undergo slower electron transfer reactions capable of downstream bioenergetic chemistry.

## Conclusions

Our design pipeline establishes that flavin excited state dynamics and redox properties can be accurately programmed within entirely *de novo* protein scaffolds, with near-atomistic precision. The 4D2 architecture supports robust dual cofactor incorporation, with crystallographic validation highlighting the accuracy now achievable in cofactor-centric protein design. Preservation of the functional heme-binding module in FC-4D2, or its selective removal in Fl-4D2, underscores the modularity of our helical bundle architectures, and their potential to encode more complex, multi-component photochemical assemblies.

A key advance is integrating quantum chemical simulation with machine-learning-driven design to deterministically target excited-state behaviour. This enabled the transformation of the fluorescent Fl-4D2 into Fl-4D2_CT_ in which the excited state landscape is deliberately reshaped. Consequentially, the electronic absorption and redox properties can be predictively modulated, with fluorescence and photoreduction quenched through ultrafast charge transfer from the flavin isoalloxazine to π-stacked tryptophan side chains, demonstrated by ultrafast spectroscopy.

More broadly, these results establish alternative routes to a wide range of operational regimes of flavin chemistry, encoding photochemical function directly from sequence by coupling data-driven design, molecular dynamics, electronic structure theory, and time-resolved spectroscopy. The ability to modulate excited-state properties within proteins opens opportunities to construct bespoke photoredox systems with pre-defined energy landscapes for targeted function. Such control will be valuable for developing light-driven catalysis, bioinspired energy conversion, and protein-based optoelectronic components, where precise management of electron flow and excited-state dynamics is essential.

## Methods

### Protein Design

All computation was performed using the University of Bristol ACRC’s High Performance Computing environments BlueCrystal Phase 4, BluePebble or Isambard-AI.^51^ Iterative design of the three-module flavocytochrome maquette was performed with LigandMPNN^20^ sequence redesign, omitting the heme-module and outputting a batch size of 250 residues on a single CPU node. A bias of –1.0 was applied to alanines due to high propensity of occurrence in previous runs, and residues histidine and cysteine omitted to prevent off-target cofactor loading. Sequences were filtered to remove duplicate sequences and those containing >3 poly-amino acid stretches, due to complexity in synthesis of homopolymeric DNA sequences.^52^ Remaining sequences were folded using ESMFold model esm.pretrained.esmfold.v1 on GPU nodes.^21^ Physicochemical properties of sequences were extracted using Biopython module Bio.SeqUtils.ProtParam^53^ and sequences ranked through a normalisation metric of LigandMPNN overall/ligand confidence and ESMFold pTM and average pLDDT. The top output was manually docked with ligands before acting as input for next iteration cycle. Fl-4D2 was designed by mutating FC-4D2 with the corresponding 9 residues that collapsed the heme site in the m1-4D2^16,18^ design process: H9L, G62W, A85L, A13I, M65L, H81F, Y131I, G134W and M137L. Single structure predictions were performed on the ESMFold_advanced Colab.^54^

The X-ray crystal structure for Fl-4D2 was used as the basis for computational redesign to tune its biophysical and photophysical properties. 14 amino acid residues surrounding the 8-demethylriboflavin (8-dMeRF) moiety were selected for redesign. LigandMPNN was used to generate 250 sequences, with a bias of –1.0 to Ala and +3.0 for all aromatic residues. The sequences were filtered as above and the top 10, ranked by overall and ligand confidence, were folded with Chai-1^34^ alongside the cysteinyl-8-dMeRF ligand, with the model selected for its accuracy and ease of incorporation of covalent modifications. The highest scoring output (pTM, ipTM) was funnelled into the consecutive round of sequence redesign and structure prediction, for a total of 3 cycles of iterative design. The highest scoring design of each iteration was used as the input for MM MD simulations.

### MM MD Simulations

The predicted protein structure was used as a starting point for classical molecular dynamics simulations under periodic boundary conditions using AMBER v24.^35^ The structures were solvated in a truncated octahedron water box of TIP3P molecules using the solvateoct plugin of Tleap in Ambertools v24.^55^ The unit cell parameters were a = b = c = 76.986 Å, α = β = γ = 109.471°, and the solvated structure was neutralised with 3 Na^+^ ions. The time step for the simulations was 2 fs, applying the SHAKE algorithm to constrain the bonds with hydrogen atoms.^56^ The non-bonded cut-off chosen was 12 Å, and the long-range interactions were handled using the Particle Mesh Ewald algorithm.^57^ The Langevin thermostat with a collision frequency of 1.0 ps^-1^ was used throughout the simulations.^58^ The Berendsen barostat was used to maintain pressure, with a pressure coupling timescale of 1.0 ps.^59^ The protein was described with the Amber ff14SB force field. Force field parameters for the riboflavin analogue, 8-dMeRF, were generated with mdgx using the same approach as our previous work.^40^

The solvated systems were energy-minimized for 1000 steps using the steepest descent method. The energy-minimised systems were heated for 1 ns until the temperature reached 300 K in the NVT ensemble. The heated systems were equilibrated for a period of 1 ns in the NPT ensemble at conditions of 300 K and 1 bar pressure. The system was considered equilibrated when the density reached 0.98 g/cm^3^. The equilibrated systems were then used as starting points for running production simulations which were carried out for 5 x 50 ns (FC,Fl-4D2) or 100 ns (Fl-4D2_CT_) in the NVT ensemble. Hydrogen bonding interactions between the 8-dMeRF and protein residues were calculated using cpptraj in Ambertools v24 for each frame within the MM trajectories.

### Computing spectra from QM/MM MD

Snapshots exhibiting parallel π-stacking interactions between 8-dMeRF and aromatic protein residues were identified from the production MM MD trajectories using a custom Python script with MDAnalysis.^60,61^ For each frame, the atoms of the isoalloxazine ring system of 8-dMeRF and the aromatic ring atoms of each protein residue were used to calculate ring centroids and best-fit ring planes. Ring centroids were defined as the geometric centre of the aromatic ring atoms, while ring-plane normal vectors were obtained by least-squares fitting of a plane to the aromatic ring atoms using singular value decomposition. π-stacking interactions were identified using three geometric criteria: (i) a centroid–centroid distance of less than 4Å, (ii) an angle of less than 30° between the ring-plane normal vectors, corresponding to approximately parallel aromatic systems, and (iii) a lateral offset of less than 2Å.^62,63^ Frames satisfying all three criteria were extracted from the production trajectories. These frames were inspected visually, and a structure exhibiting a π-stacking geometry was selected as a starting conformation for QM/MM molecular dynamics simulations. For Fl-4D2_CT_, the snapshot selected met the above criteria with the following geometric quantities: (i) 3.58 Å, (ii) 14.91°, (iii) 1.39 Å.

QM/MM MD was performed with Amber v24/TeraChem v1.9-2023.06-dev.^36–39^ The QM region for QM/MM MD was 8-dMeRF described using DFT at the wB97X-D3/def2-SVP level of theory, while the surrounding protein and solvent was included in the MM region described with the Amber forcefield ff14SB and TIP3P respectively. The interaction between the QM and MM regions was described using electrostatic embedding with a point charge cutoff of 12 Å. 10 ps dynamics were conducted in the NVT ensemble. The system reached 300 K after 2 ps for the whole system, as well as the composite QM and MM regions. 100 snapshots, sampled every 0.08 ps, were extracted between 2 ps and 10 ps of the QM/MM trajectory for further processing. 15 excited states were calculated for each snapshot with linear-response time-dependent density functional theory (LR-TDDFT) at the same level of theory, employing the Tamm-Dancoff approximation (TDA).^64^ Residues with atoms within 3 Å of 8-dMeRF were included in the QM region for LR-TDDFT calculations to more accurately account for the effect of the protein environment on 8-dMeRF’s absorption spectrum. The excitation energies and transition dipole moments were extracted, and the spectrum was then broadened with a Gaussian function with a FWHM of 0.165 eV.^65^

The natural transition orbitals (NTOs) were calculated using TheoDORE 2.4.20^66^ and visualised in ChemCraft.^67^ The charge-transfer (CT) character between 8-dMeRF and surrounding residues was calculated using TheoDORE v2.4.20. The fragments input into this analysis were the atoms of the 8-dMeRF isoalloxazine ring and the atoms of the closest aromatic residues. These were Phe27 and Phe99 for R1 and Trp27 and Trp99 for Fl-4D2_CT_. For each snapshot, a CT number was calculated between the 8-dMeRF and each aromatic residue for all 15 electronic transitions. As either aromatic residue could act as the dominant CT partner depending on the geometry, only the larger of the two residue-specific CT contributions per snapshot was used for the final (max) CT number averages. The mean of all these values were calculated over all 100 snapshots.

### Protein Expression and Purification

Designed sequences were translated to DNA and appended downstream of a ribosome-binding site, His_6_-tag, V5 epitope and TEV protease cleavage site (Fig.S3), before being ordered pre-cloned into pET21(+) vectors (TWIST Bioscience) and transformed into T7 Express *E. coli* cells (NEB). Transformation colonies were used to inoculate 100 mL LB cultures containing 50 µg/mL carbenicillin which were incubated overnight at 37 °C, 200 RPM. 20 mL overnight cultures were added to 1 L 2x YT media in 2 L baffled flasks alongside carbenicillin and grown at 37 °C, 200 RPM until an OD_600_ of 0.7-0.8, of which they were then induced with 1 mM isopropyl β-D-1-thiogalactopyranoside (IPTG) for 4 h. Cells were harvested at 4392 x*g*, 4 °C, 20 mins and resuspended in lysis buffer (50 mM monobasic sodium phosphate, 300 mM sodium chloride, 20 mM imidazole, and 1 mM tris(2-carboxyethyl)phosphine (TCEP), pH 8.0).

The cells were disrupted through sonication (FisherBrand Model 120 Sonic Dismembranator, 6 mm probe, 20 s on 20 s off, 80% amplitude) before further centrifugation at 38,759 x*g*, 4 °C, 35 mins. Cell lysates were loaded onto HisTrap FF Ni-NTA columns (Cytiva) equilibrated in lysis buffer, with bound proteins eluted by raising the imidazole concentration to 250 mM. Proteins were then exchanged into denaturing buffer (6 M guanidine hydrochloride, 8 mM imidazole, 50 mM monobasic sodium phosphate, 50 mM Tris(hydroxymethyl)aminomethane (TRIS), 1 mM TCEP, pH 9.0) with 7-methyl-8-chloro-10-(1’-D-ribityl)isoalloxazine (8-chloro-8-dMeRF, see Section S4) solubilised to 6 mg/mL in dimethylformamide added in a 2x molar excess, before overnight incubation at 50 °C, 200 RPM. Excess flavin was separated by flavin-loaded His_6_-tagged protein with an additional Ni-NTA chromatography as above. The protein was then exchanged into 50 mM TRIS, 0.5 mM EDTA, 1 mM TCEP, pH 8.0 and hTEV60 protease^68^ added in a 1:30 protease:protein ratio. After overnight incubation at 30 °C, 180 RPM, the His_6_-cleaved protein was purified with a further Ni-NTA affinity chromatography before size exclusion chromatography (SEC) using a HiLoad 16/600 Superdex 75pg column (Cytiva) equilibrated in 20 mM 2-(cyclohexylamino)ethanesulfonic acid (CHES), 100 mM KCl at pH 8.6.

As any endogenous heme was lost upon denaturation, FC-4D2 was then supplemented with a 1.5x molar excess of hemin from a 10 mg/mL stock in dimethyl sulfoxide (DMSO). The mixture was incubated overnight on rollers at room temperature before excess heme removed using PD-10 desalting columns packed with Sephadex G-25 resin. An additional round of SEC was performed to yield the final purified product.

Different pHs were assessed in the photophysical analysis, with protein samples exchanged using desalting columns into buffers of either pH 6.5 (20 mM 2-(N-morpholino)ethanesulfonic acid (MES), 100 mM KCl) or pH 7.6 (20 mM 4-(2-hydroxyethyl)-1-piperazineethanesulfonic acid (HEPES), 100 mM KCl).

### Protein Crystallography

Protein crystallisation of FC-4D2 and Fl-4D2 was screened using the sitting drop method of vapour diffusion. MRC 96-well 2-Drop plates were loaded with commercially available screens (Molecular Dimensions) and protein samples of 12 mg/mL in 20 mM CHES, 100 mM KCl, pH 8.6 dispensed in ratios of 150:150 (nL) and 125:175 (nL) protein:screen. Trays were incubated at 20 °C and crystals were well-formed within a week. Diffraction quality crystals of FC-4D2 grew in 2.0 M ammonium sulfate, 0.1 M bis-tris, pH 5.5, and of Fl-4D2 in 0.09 M Halogens [0.03 M sodium fluoride, 0.03 M sodium bromide, 0.03 M sodium iodide], 0.1 M Buffer System 1 [0.06 M MES, 0.04 M imidazole], 37.5% v/v Precipitant Mix 4 [9.4% v/v MPD, 9.4% w/v PEG 1000, 9.4 w/v PEG 3350], pH 6.5. Crystals selected for data collection were mounted in LithoLoops (Molecular Dimensions) and flash frozen in liquid nitrogen. Diffraction data were collected at Diamond Light Source, UK, on beamlines i04 and i24. Data were autoprocessed, merged and scaled by Xia2^69^ using the Diamond Light Source Automatic Software Pipeline. The structures were determined by molecular replacement using MOLREP^70^ with the ESMFold^21^ predicted structures of the proteins as search models. Iterative rounds of structure refinement was performed using the CCP4i2^71^ software suite tools COOT^72^ and Refmac5^73^. Protein structure images were prepared using PyMOL 3.1.^74^

### Protein Film Cyclic Voltammetry

Working electrodes were comprised of mesoporous indium-tin oxide (20 µm thickness, 0.25 cm^2^ footprint (geometrical surface area), and 750 nm pore diameter) on fluoride-doped tin oxide–coated glass following the protocol presented by Mersch *et al*.^75^ Solutions containing 100–200 µM protein (20 mM CHES, 100 mM KCl, pH 8.6) were drop cast onto ice-cold electrodes and left for 15 min before transfer to a N_2_ chamber (atmosphere < 10 ppm O_2_) where they were rinsed with the same buffer-electrolyte solution to remove loosely bound protein. The working electrode was then introduced to a three-electrode cell located inside a Faraday cage within the N_2_ chamber. That cell had a Pt counter electrode, Ag/AgCl (saturated KCl) reference electrode and contained 20 mM CHES, 100 mM KCl, pH 8.6. Cyclic voltammetry was performed with a PGSTAT30 potentiostat (Metrohm Autolab) under the control of NOVA software.

Faradaic (protein associated) currents were isolated from the measured data used QSoaS^76^ and measured potentials converted to values against standard hydrogen electrode (SHE) by the addition of 0.195 V. Contributions to the Faradaic current from Fl-4D2 and Fl-4D2_CT_ were then modelled in Excel using Eq. 1^77^ to describe the current *i(E)* as a function of potential *(E)* from a flavin cofactor having reduction potentials *E_1_* and *E_2_*, describing the ox/semi-quinone and semi-Q/red couples respectively.

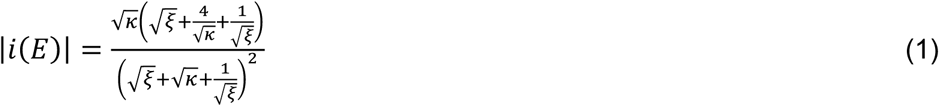

Here, κ = exp(*F*(*E_1_ – E_2_*)*/RT*) and ξ = exp(*-2F*(*E – E_m_*)/*RT*), with *F*, *R*, *T* having their usual meanings and *E_m_* being the flavin midpoint potential for the modelled peak, (*E_1_ + E_2_*)/2. For FC-4D2 which contains heme in addition to flavin, the Faradaic currents were modelled as the sum of contributions from flavin Eq. 1 and heme Eq. 2 assuming 1:1 heme:flavin binding within FC-4D2 such that the area of the modelled flavin peak was twice that of the modelled heme peak.

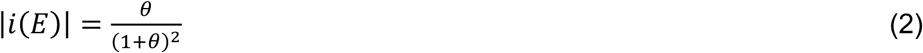

Whereby θ = exp(*F*(*E* – *E_m_*)/*RT*), with *E_m_* representing the single heme 3+/2+ midpoint potential. For each cofactor, the *E_1_/E_2_/E_m_* values reported are averages obtained from modelling of the corresponding oxidation and reduction peaks, collected from at least one scan of three individually produced electrodes.

Fit quality was calculated as one minus the mean absolute deviation between the modelled (*i_mod_*) and experimental currents (*i_exp_*), normalised to the peak current (*i_p_*) and expressed as a percentage, Eq. 3. Here *E_x_* represents each potential value accessed and *N* is the total number of data points in the cyclic voltammogram.

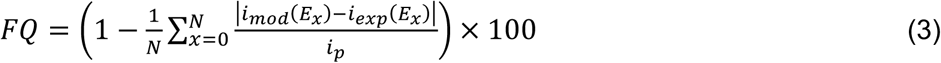

### Continuous Wave Photoreduction

Photoreduction data was collected upon continuous wave (cw) white-light irradiation (150 W xenon lamp) using a SX20 system (AppliedPhotophysics) stopped-flow apparatus under semi-anaerobic conditions ([O_2_] < 2 ppm). The fluence of the cw white light (∼400-700 nm) on the sample was 7.85 × 10^−4^ J s^−1^ mm^−2^. Deoxygenated samples were prepared to an OD at λ_max_ of roughly 0.6-0.7 and shot against equal volumes of buffer and spectra recorded with a multiwavelength photodiode array detector. As complete photoreduction was not achieved upon cw irradiation, spectra was also collected upon addition of molar excess of reducing agent sodium dithionite, in order to obtain a spectrum of the fully reduced species. The spectra were deconvolved into linear contributions of each ligand redox state and the dithionite itself where applicable, with extinction coefficients of 8043 M^−1^cm^−1^ at 315 nm used for sodium dithionite^78^, and ∼12300 M^−1^cm^−1^ for the 8-dMeRF due to the concentration not being known precisely.^79^ The basis spectra for a single bis-histidine heme B for FC-4D2 was derived from m1-m4D2.^16,18^

### Ultrafast Transient Absorption Spectroscopy

Femtosecond to microsecond transient absorption spectroscopy measurements were carried out using an established dual synchronised amplifier laser system (2x Pharos PH2, Light Conversion).^40^ The primary laser amplifier, operating at 1 kHz, drove an optical parametric amplifier (Orpheus-HP, Light Conversion) to generate pump laser pulses (480 nm, FWHM = 4 nm). The secondary amplifier, operating at 2 kHz, was used to generate a white light supercontinuum probe pulse (spanning 340–850 nm). A small portion of the 1030 nm fundamental was focused into a continually rastered CaF_2_ crystal (Crystran, 5 mm thick). The residual fundamental near-infrared light was removed from the probe beam using a Schott KG3 glass filter.

The protein samples were prepared in a semi-anaerobic chamber to an OD of 0.6 at the excitation wavelength with 20 mM EDTA and loaded in cuvettes of 1 mm pathlength (Starna Scientific, 1/GL14/SOG/1). At the sample position, the pump was focussed to a spot size of 160 µm diameter with pulse energy of 300 nJ/pulse (12.4 GW cm^−2^). The overlapping probe pulse was focused to a spot size of 60 µm. To avoid sample bleaching, the sample cuvette was rastered in plane. The signal and probe beams were collimated and imaged into a homebuilt spectrograph, coupled to a Glaz LineScan-I-Gen2 camera (Synertronic Designs). Data were acquired using reference detection (*via* simultaneous acquisition of a reference probe beam that did not pass through the sample) to account for shot-to-shot intensity fluctuations in the white-light generation. The polarization of the probe was set to magic angle with respect to the pump using a half-waveplate-polariser pair (WPH10M-1030 and WP25M-UB1, Thorlabs). The instrument response was measured to be 120 fs *via* the non-resonant response from buffer solution.

### Pulsed Photoreduction

Pulsed photoreduction data were acquired using the same 480 nm laser source as in TA experiments, but at a higher repetition rate of 10 kHz. The entire sample, contained in a 1 mm cuvette, was irradiated with a laser fluence of 4.2 × 10^−4^ J s^−1^ mm^−2^ for time increments increasing from 20 s. This corresponded to a peak pulse power intensity of 2.4 MW cm^−2^. After each irradiation increment the steady-state absorption was measured on a Cary 60 UV/Visible Spectrophotometer (Agilent Technologies). To probe the effect of exciting red-absorbing photoproducts (*e.g.* the semiquinone), photoreduction was also repeated by simultaneously irradiating with the pulsed 480 nm laser and an additional continuous-wave narrowband HeNe 632.8 nm laser (98 × 10^−6^ J s^−1^ mm^−2^).

## Data Availability

X-ray crystallographic structures have been deposited at the PDB under 30AK and 29RO. All (QM) MM MD trajectories are contained within the University of Bristol data repository (DOI: 10.5523/bris.aajxpbpjac7g24rb814q21knj).

## Author Contributions

The study was conceived by E.L.B., J.U., A.J.M., T.A.A.O. and J.L.R.A. E.L.B. conducted the *in silico* design of FC- and Fl-4D2, as well as electrochemical and biophysical characterisation of each design. J.U. led the QM/MM-directed protein design with guidance from A.G.R. and B.F.E.C. and conducted the biophysical characterisation of Fl-4D2_CT_ with E.L.B. A.N.S. and M.L. led time-resolved spectroscopic studies. X-ray diffraction data was collected by C.R.B and R.B., with molecular replacement and refinement performed by C.R.B, R.B., P.H. and E.L.B. Organic synthesis of the flavin analogue was performed by J.C.B-R. and B.R.L. NMR data was collected and analysed by P.D. and C.W. J.A.B. collected and analysed the RFQ EPR data. Collection and analysis of PFE data was performed by E.L.B, under guidance from J.N.B. All authors contributed to manuscript writing.

## Acknowledgements

This work was supported at the University of Bristol, University of Portsmouth and University of East Anglia by the Biological and Biotechnological Sciences Research Council (BB/W003449/1). E.L.B., J.U. and M.L. acknowledge PhD funding from University of Bristol scholarship, EPSRC doctoral training award, and Colin Western Bequeath, respectively. B.F.E.C. thanks EPSRC for funding through the Programme Grants EP/V026690/1 (UPDICE) and EP/X026973/1 (COSMOS). A.G.R. and P.H. acknowledge funding from the ERC (grant agreement No. 101021207). We would like to thank Dr Dima Svistunenko for his assistance in rapid-freeze quench sample preparation. We would also like to thank Alexander L. Sutton-Cook for the preparation of ITO working electrodes for PFE. The authors would like to thank Diamond Light Source for beamtime (proposals mx31440 and mx37593), and the staff of beamlines I04 and I24. The authors acknowledge the use of resources provided by the Isambard-AI National AI Research Resource (AIRR). Isambard-AI is operated by the University of Bristol and is funded by the UK Government’s Department for Science, Innovation and Technology (DSIT) *via* UK Research and Innovation; and the Science and Technology Facilities Council [ST/AIRR/I-A-I/1023]. The authors also acknowledge the Advanced Computing Research Centre at the University of Bristol for High-Performance Computing resources, particularly the facilities and staff on clusters BlueCrystal Phase4 and BluePebble.

