## Supporting Information for "*De Novo* Design of Flavoproteins with Controlled Excited State Dynamics"

### S1 Design and biophysical characterisation

## S1.1 FC-4D2 & FI-4D2

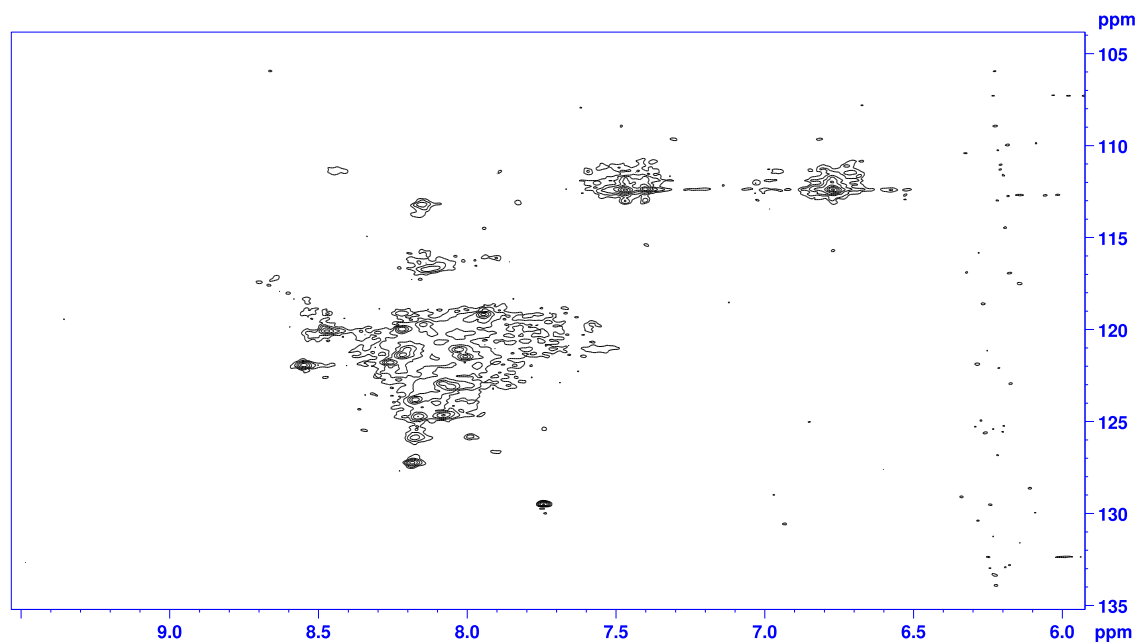

**Supplementary Figure S1:** $^1\text{H}$-$^{15}\text{N}$ SOFAST-HMQC spectra of initial design termed FI-4D2\_cov5.<sup>1</sup> A lack of peak dispersity and resolution indicated poor structural homogeneity when the Rosetta-designed scaffold was bound to heme B and 8-demethylriboflavin (8-dMeRF). This prompted the evaluation of a longer, three-module scaffold with a central hydrophobic packing domain to encourage more consistent, thermostable folding.

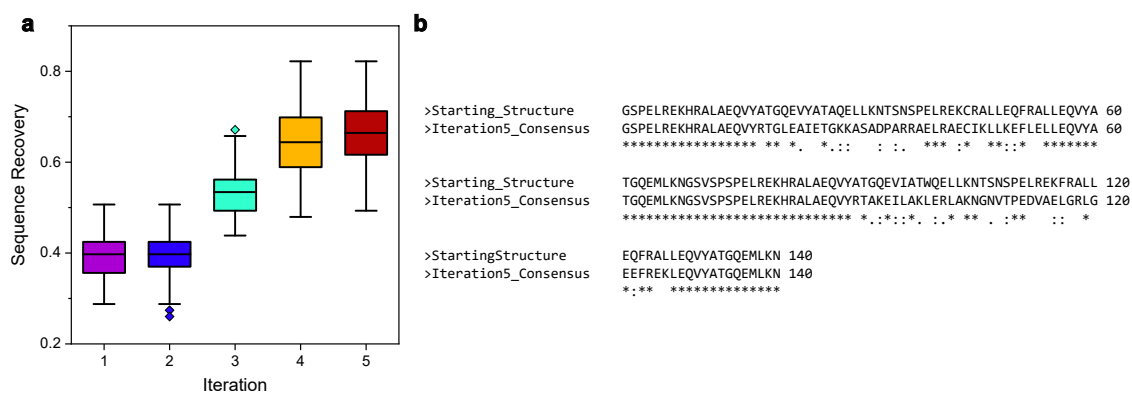

**Supplementary Figure S2:** Sequence convergence in the iterative design of FC-4D2. (a) Sequence recovery per design iteration. (b) Iteration 5 consensus sequence against the starting structure constructed from m2-4D2, demonstrating the sequence diversification of the flavin binding module.

All constructs have an N-terminal His<sub>6</sub>-tag, V5 epitope and TEV protease cleavage site:

Amino acid (AA) sequence: MHHHHHHGKPIPNPLLGLDSTENLYFQ

DNA sequence: ATGCATCATCACCATCACCATGGTAAGCCTATCCCTAACCCTCTCCTCGGTCTCGATTCTACGGAAAACCTGTATTTTCAG

#### >FC-4D2

AA GSPELREKHRALAEQVYRTGLRAIETGERASADPARREALRAECIRLLEEFLRLLEQVYATG
QEMLKNGSVSPSPELREKHRALAEQVYETAKEILAKLKRLAADGRVTPEDVAELGRLGEEFR
ENLEQVYATGQEMLKN

DNA GGGTCTCCAGAACTTCGCGAAAAGCACCGGGCGCTGGCCGAACAAGTCTATCGTACTG
GACTGCGTGCCATTGAAACGGGCGAACGCGCGTCGGCGGATCCGGCACGCCGCGAGG
CTTTAAGAGCGGAGTGCATCCGTCTCCTCGAGGAATTTCTGCGTCTGTTAGAACAGGTG
TATGCCACCGGTCAGGAGATGCTGAAAAACGGTAGCGTTTCCCCTAGTCCCGAACTTCG
GGAAAAGCATCGCGCATTAGCGGAACAGGTATACGAAACAGCGAAAGAGATTCTGGCT
AAATTGAAACGACTGGCCGCAGATGGTCGTGTGACGCCGGAAGACGTTGCAGAATTGG
GCCGTCTAGGCGAAGAGTTCCGCGAAAATCTGGAGCAAGTGTACGCTACCGGCCAGGA
AATGCTGAAAAACTAA

#### >FI-4D2

AA GSPELREKLRALIEQVYRTGLRAIETGERASADPARREALRAECIRLLEEFLRLLEQVYATWQ
ELLKNGSVSPSPELREKFRALLEQVYETAKEILAKLKRLAADGRVTPEDVAELGRLGEEFRE
NLEQVIATWQELLKN

DNA GGTTCGCCGGAATTACGTGAAAAGCTGCGTGCGCTTATCGAGCAAGTGTATCGCACGG
GCCTGCGTGCCATCGAAACTGGCGAGCGTGCATCCGCCGATCCGGCACGCAGAGAGG
CCTTGCGTGCTGAATGCATTCGCCTGCTCGAAGAATTTCTTCGGCTGCTGGAGCAGGTG
TATGCTACCTGGCAGGAACTGTTGAAAAACGGGAGTGTCAGCCCATCTCCCGAACTTCG
CGAAAAATTCCGGGCGTTGCTGGAGCAGGTTTACGAGACCGCGAAAGAAATTTTAGCAA
AACTGAAGCGTCTGGCGGCTGACGGTCGCGTTACGCCTGAGGATGTAGCGGAACTCGG
CCGCCTGGGAGAAGAATTTCGAGAAAATTTAGAACAAGTGATTGCCACATGGCAGGAAC
TACTGAAAAACTAA

###### >FI-4D2<sub>CT</sub>

AA GSPELREKLRALIEQVYRTGLRAEEAWRRARADPAREEALRAECIRLLEEFLRLLEQVYATW
QELLKNGAASPSPELREKFRALLEQVYETAKELLAKWKRIAADGRLTPEDEAELRRLGEEFR
ENLEQVIATWQELLKN

DNA GGTAGTCCGGAGCTGCGCGAGAAATTGCGTGCGCTTATTGAACAGGTATACCGCACCG
GTTTACGTGCAGAAGAAGCCTGGCGCCGTGCGCGGGCCGATCCGGCGCGTGAAGAAG
CGCTGAGAGCAGAATGCATTCGCCTCTTAGAGGAATTTCTGCGCCTACTGGAACAGGTC
TATGCCACTTGGCAAGAACTGCTCAAAAACGGCGCTGCATCCCCAAGCCCCGAACTGC
GTGAAAAATTCCGGGCCCTGCTTGAGCAGGTGTATGAAACAGCTAAGGAATTACTGGCG
AAATGGAAGCGTATCGCGGCTGATGGGCGACTGACGCCTGAGGACGAGGCCGAGCTG
CGCCGCCTGGGCGAAGAATTTAGGGAAAATTTGGAGCAGGTTATCGCAACCTGGCAAG
AACTTTTGAAAAACTAA

**Supplementary Figure S3:** DNA and amino acid sequences of FC-4D2, Fl-4D2 and FI-4D2<sub>CT</sub>.

|  |  |  |
| --- | --- | --- |
| F1-4D2CT | GSPELREKLRALIEQVYRTGLRAEEAWRRARADPAREEALRAECIRLLEEFLRLLEQVYATWQELLKNGA | 70 |
| FC-4D2 | GSPELREKHRALAEQVYRTGLRAIETGERASADPARREALRAECIRLLEEFLRLLEQVYATGQEMLKNGS | 70 |
| F1-4D2 | GSPELREKLRALIEQVYRTGLRAIETGERASADPARREALRAECIRLLEEFLRLLEQVYATWQELLKNGS | 70 |
|  | ***** ** |  |
| F1-4D2CT | ASPSPELREKFRALLEQVYETAKELLAKWKRIAADGRLTPEDEAELRRLGEEFRENLEQVIATWQELLKN | 140 |
| FC-4D2 | VSPSPELREKHRALAEQVYETAKEILAKLKRLAADGRVTPEDVAELGRLGEEFRENLEQVYATGQEMLKN | 140 |
| F1-4D2 | VSPSPELREKFRALLEQVYETAKEILAKLKRLAADGRVTPEDVAELGRLGEEFRENLEQVIATWQELLKN | 140 |
|  | .*****.*** *****:*** **:*****:*** ** ***** ** **: |  |

**Supplementary Figure S4:** Multiple sequence alignment of the designed constructs.

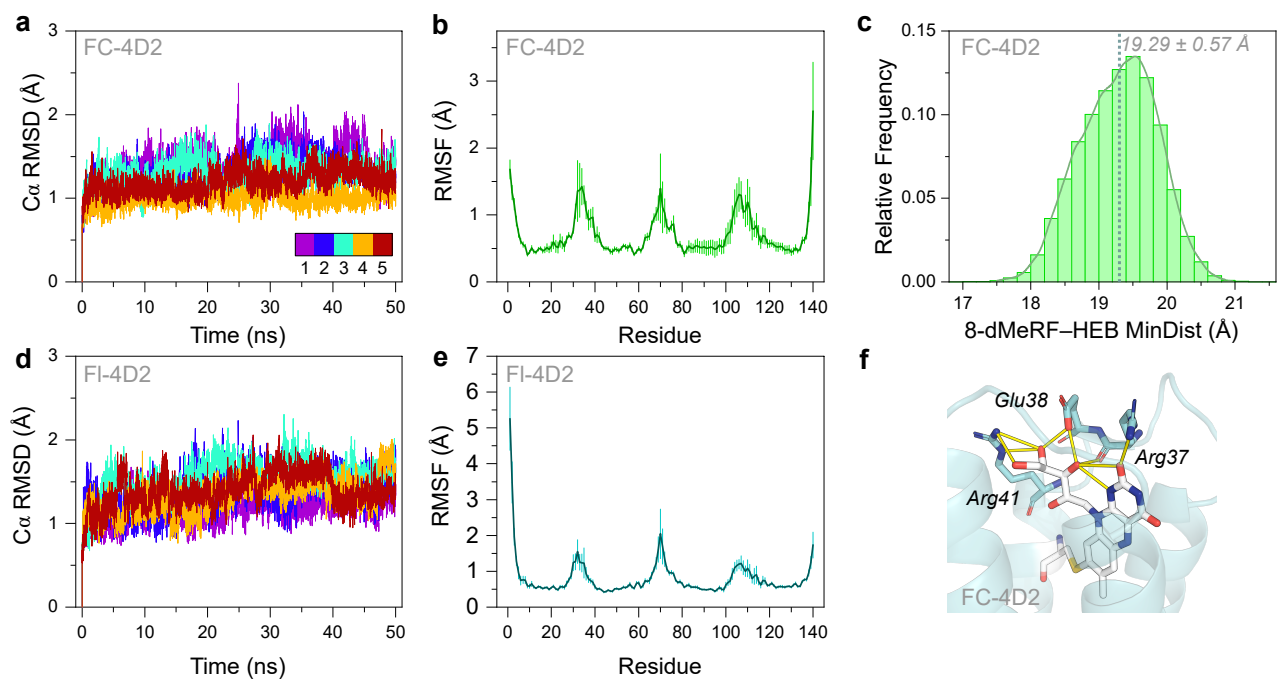

**Supplementary Figure S5:** MD analysis of FC-4D2 and FI-4D2. $C_{\alpha}$ RMSD plots for FC- (a) and Fl-4D2 (d), repeats coloured by iteration presented in colourbar. (b,e) Average heavy atom RMSF plots across 5x 50 ns for FC-4D2 and Fl-4D2 respectively. (c) Distribution plot of the minimum distance between cysteinyl linked 8-dMeRF and heme B in FC-4D2 across 5 x 50 ns MD simulation. (f) Final frame snapshot of FC-4D2 with predicted 8-dMeRF hydrogen bonding partners Arg37, Glu38 and Arg41 presented by sticks. Polar contacts as predicted by PyMOL presented as yellow lines.

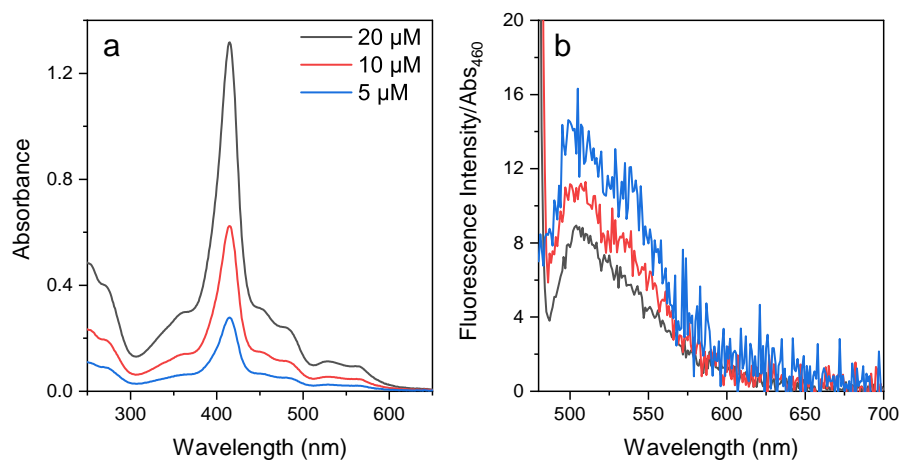

**Supplementary Figure S6:** Reducing the concentration of FC-4D2 (a) yields an increase in absorbance-normalised flavin fluorescence emission intensity (b) upon excitation with 460 nm light. This suggests that the reduced flavin fluorescence of the FC-4D2 is partly due to absorption of the excitation/emission light by the adjacent heme moiety, an example of the inner-filter effect.

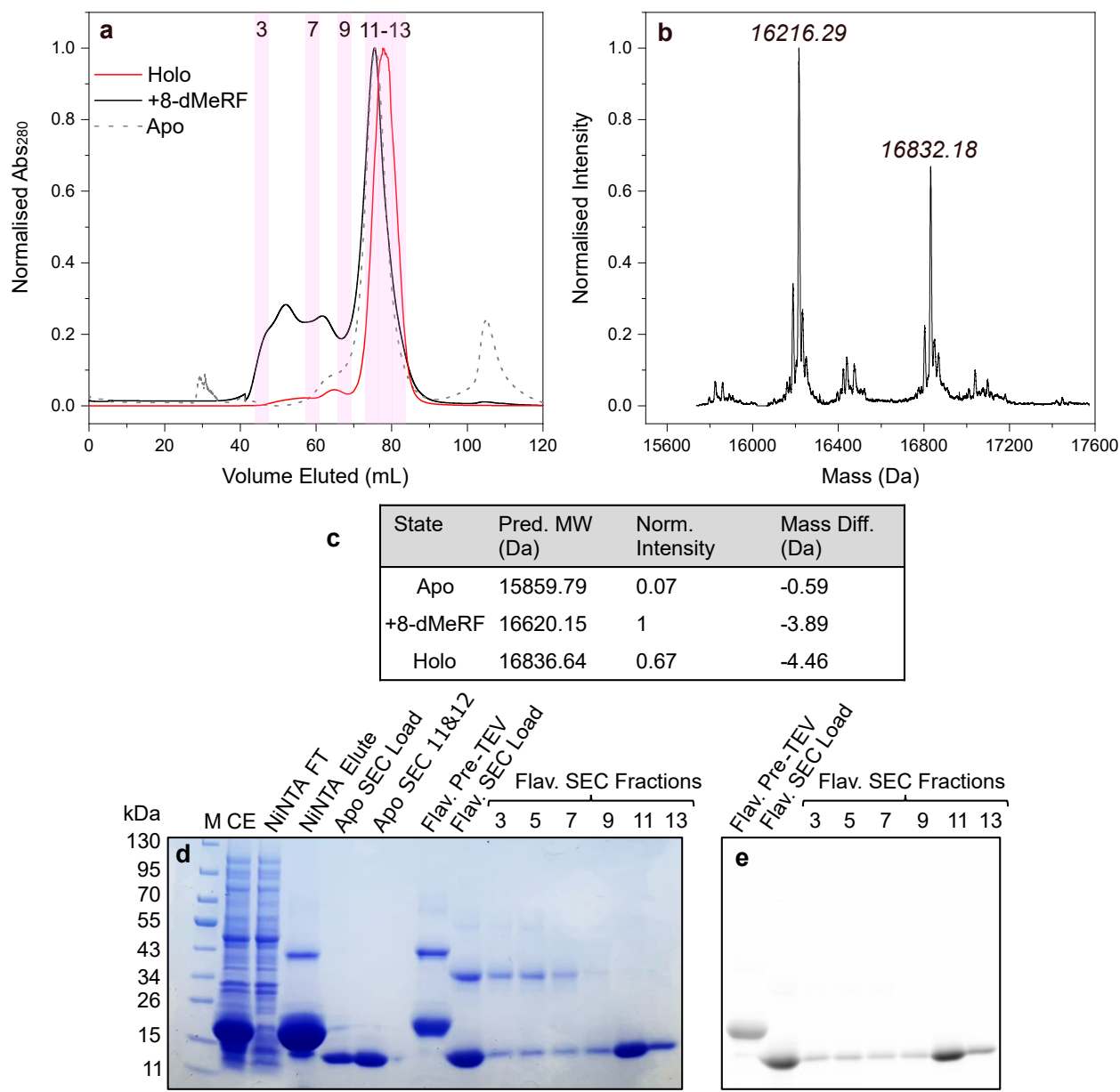

**Supplementary Figure S7:** Analytical characterisation of FC-4D2. (a) SEC chromatograms of FC-4D2 in the apo (grey dash), flavin-only ('+8-dMeRF', black) and fully-holo (red) states. Highlighted columns represent fractions analysed in SDS-PAGE. (b) Deconvoluted ESI-MS spectrum of holo FC-4D2, with masses of two prominent species indicated. (c) Table of predicted mass of each FC-4D2 state, with normalised intensity and mass difference relative to the corresponding ESI-MS peak. (d) SDS-PAGE of FC-4D2 throughout purification, with the corresponding fluorescent image of the same gel upon excitation with 473 nm laser (e).

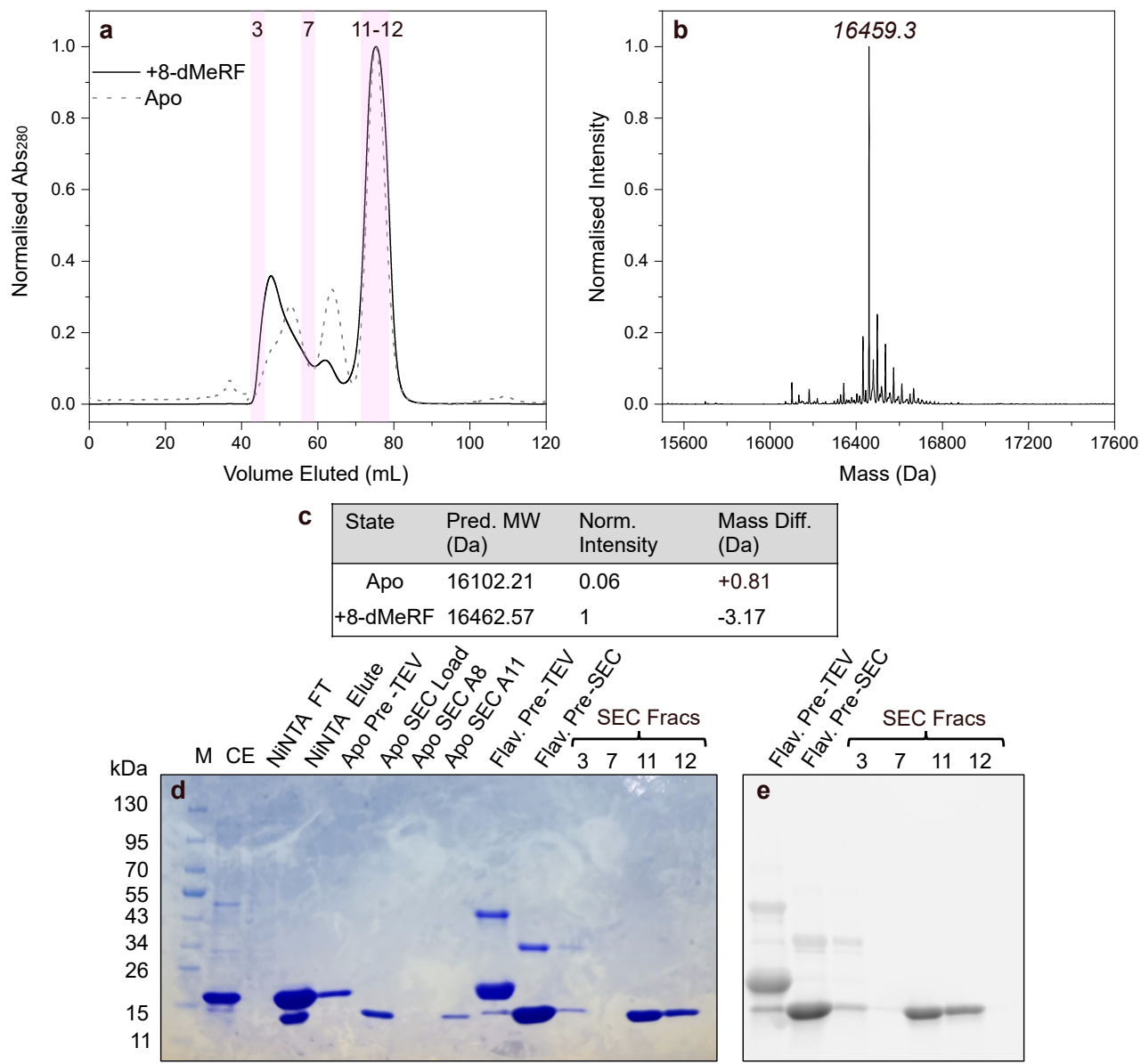

**Supplementary Figure S8:** Analytical characterisation of Fl-4D2. (a) SEC chromatograms of FI-4D2 in the apo (grey dash) and holo state ('+8-dMeRF', black). Highlighted columns represent fractions analysed in SDS-PAGE. (b) Deconvoluted ESI-MS spectrum of holo Fl-4D2, with the mass of the most prominent species indicated. (c) Table of predicted mass of each Fl-4D2 state, with normalised intensity and mass difference relative to the corresponding ESI-MS peak. (d) SDS-PAGE of Fl-4D2 throughout purification, with the corresponding fluorescent image of the same gel upon excitation with 473 nm laser (e).

|  | FC-4D2 | FI-4D2 |
| --- | --- | --- |
| **Data collection** |  |  |
| Beamline wavelength (Å) | 0.6199 | 0.9999 |
| Space group | $P2_12_12_1$ | $P12_11$ |
| Cell dimensions |  |  |
| a, b, c (Å) | 27.35, 63.02, 70.29 | 25.76, 63.44, 34.39 |
| $\alpha, \beta, \gamma$ (°) | 90.0, 90.0, 90.0 | 90.0, 92.7, 90.0 |
| Resolution (Å) | 35.14-1.35 (1.37-1.35)<sup>a</sup> | 63.44-1.8 (1.84-1.8)<sup>a</sup> |
| $R_{pim}$ | 0.026 (0.345)<sup>a</sup> | 0.028 (0.127)<sup>a</sup> |
| No. of unique reflections | 27190 (1187)<sup>a</sup> | 19110 (1073)<sup>a</sup> |
| $I/\sigma I$ | 15.0 (2.0)<sup>a</sup> | 14.3 (1.4)<sup>a</sup> |
| CC<sub>1/2</sub> | 0.998 (0.670)<sup>a</sup> | 0.998 (0.935)<sup>a</sup> |
| Completeness (%) | 98.7 (87.8)<sup>a</sup> | 99.5 (93.2)<sup>a</sup> |
| Redundancy | 10.2 (4.6)<sup>a</sup> | 1.9 (1.9)<sup>a</sup> |
| **Refinement** |  |  |
| $R_{work}/R_{free}$ | 0.154/0.188 | 0.175/0.227 |
| No. of atoms |  |  |
| Protein | 1090 | 1120 |
| Ligand/ion | 111 | 31 |
| Water | 71 | 20 |
| Average B factors (Å<sup>2</sup>) |  |  |
| Protein | 18.5 | 32.5 |
| Ligand/ion | 19.3 | 31.0 |
| Water | 30.0 | 34.8 |
| Root mean square deviations |  |  |
| Bond lengths (Å) | 0.012 | 0.007 |
| Bond angles (°) | 2.02 | 1.55 |
| Ramachandran favoured (%) | 97.7 | 97.1 |
| Ramachandran outliers (%) | 0.75 | 0.0 |

**Supplementary Table S1:** X-ray collection and structural refinement statistics. [a] values in parentheses are for the highest resolution shell.

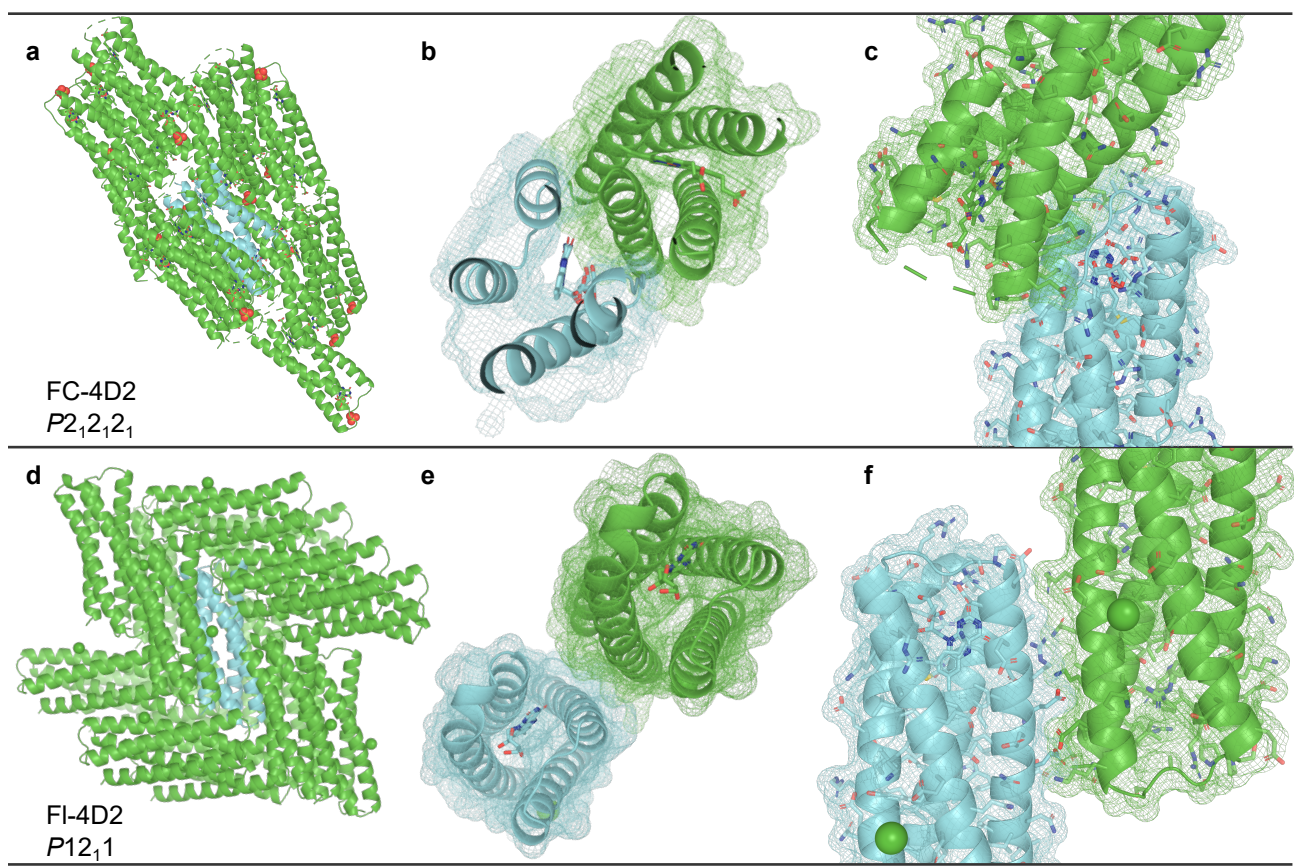

**Supplementary Figure S9:** Packing geometries in crystals of FC-4D2 and Fl-4D2. (a) Symmetry partners of FC-4D2 crystal structure in spacegroup $P2_12_12_1$ within a 12 Å radius of central cyan monomer. (b-c) Mesh surface representations of symmetry mates shows steric hindrance between Van der Waals radii of monomers, inducing a loss-of-helicity in residues 28-32 located primarily in the flavin module. (d) Symmetry partners of FI-4D2 in $P12_11$ spacegroup. (e-f) Equivalent representation of (b-c) showing an absence of clash in the flavin module of Fl-4D2 monomers, with contacts between symmetry mates occurring at discrete locations along the long-axis of each four-helix bundle.

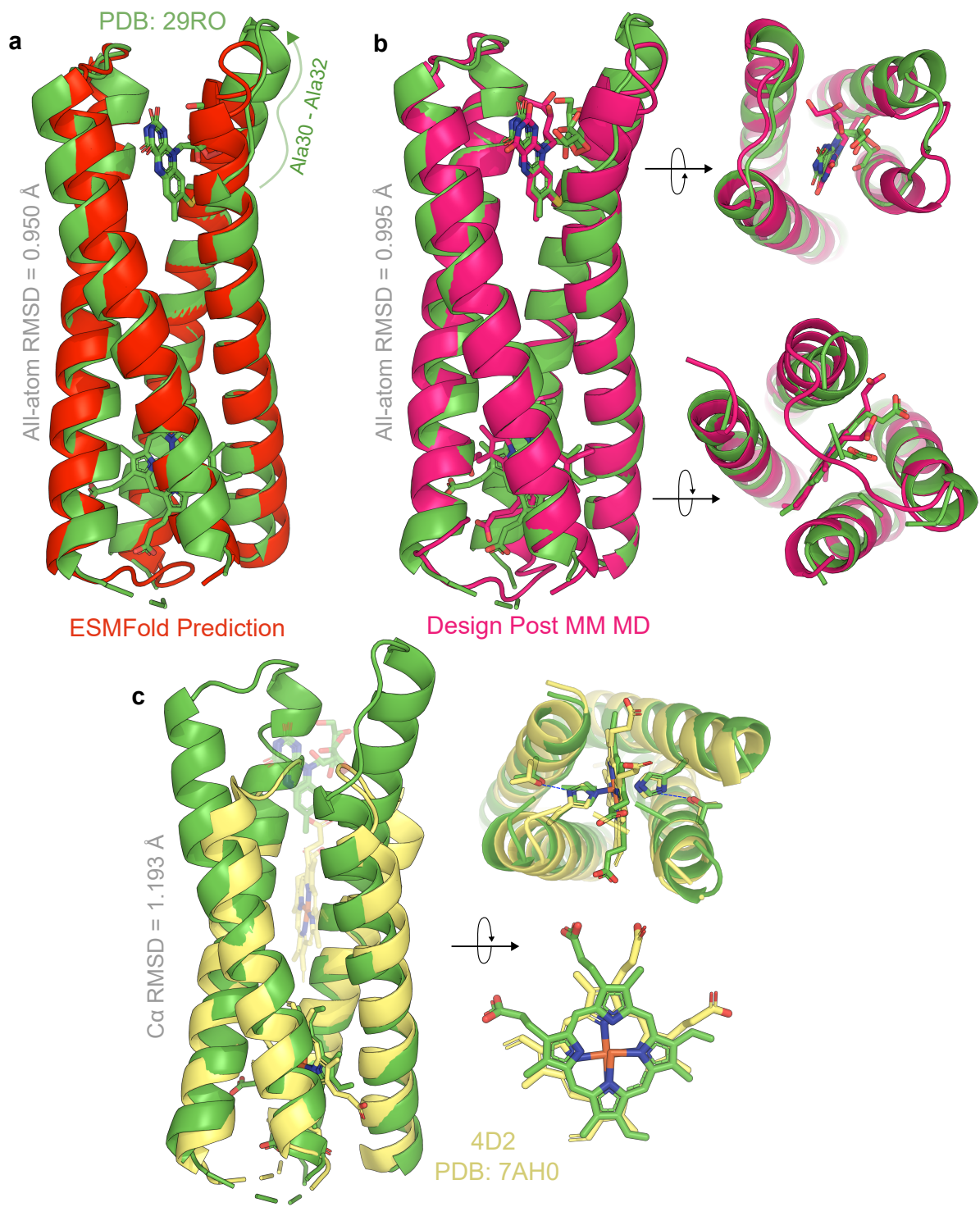

**Supplementary Figure S10:** FC-4D2 crystal structure (green) aligned with apo ESMFold model (a, red) and a final frame MM MD holo structure (b, magenta), the latter displaying top-down alignments between flavin modules, and bottom-up alignments of the heme sites. (c) Cα RMSD alignment between FC-4D2 and 4D2, demonstrating structural homology in the heme domain. The keystone secondary-coordination-sphere interactions between threonines and heme-coordinating histidines is preserved between designs (blue dashed lines), but the FC-4D2 heme sits in a 90° rotation compared to its parent protein.

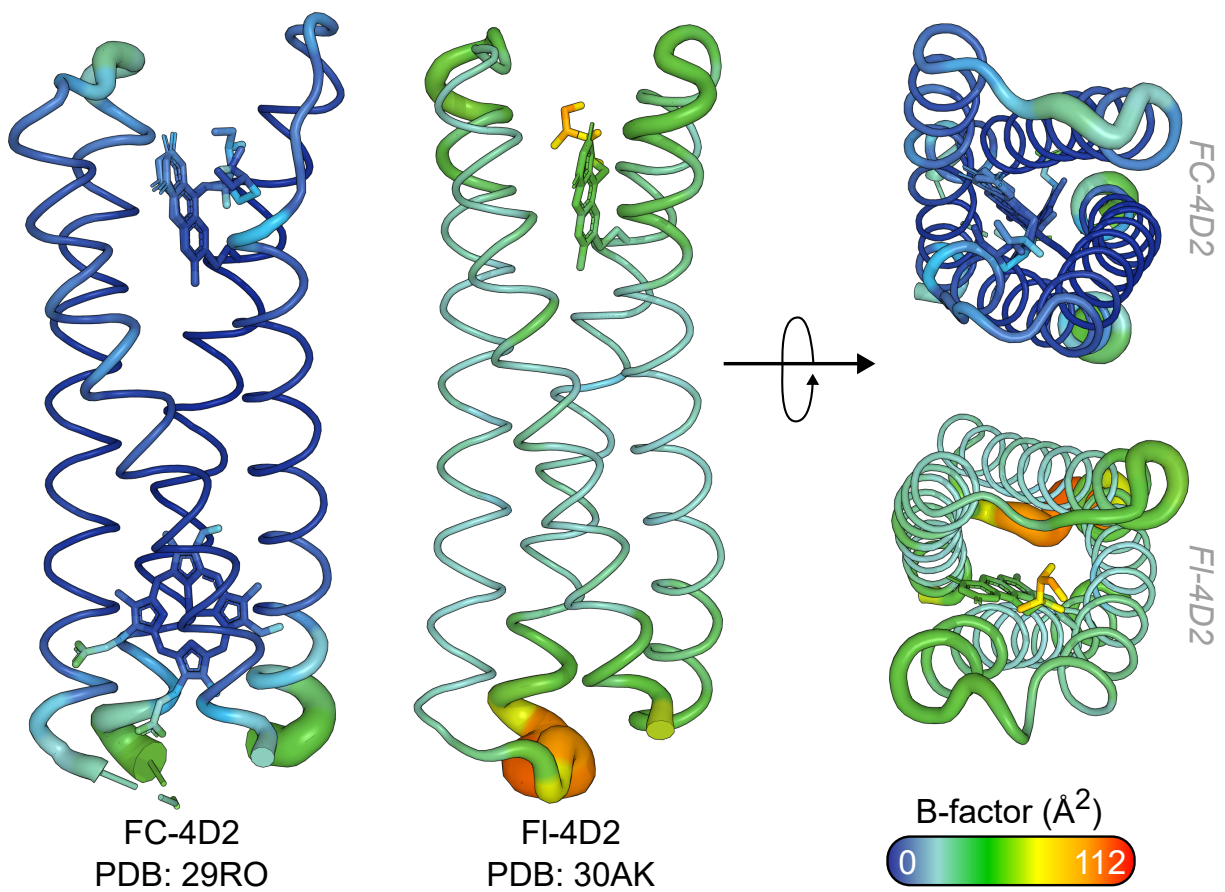

**Supplementary Figure S11:** Crystallographic B-factors mapped onto the structures of FC-4D2 and FI-4D2.

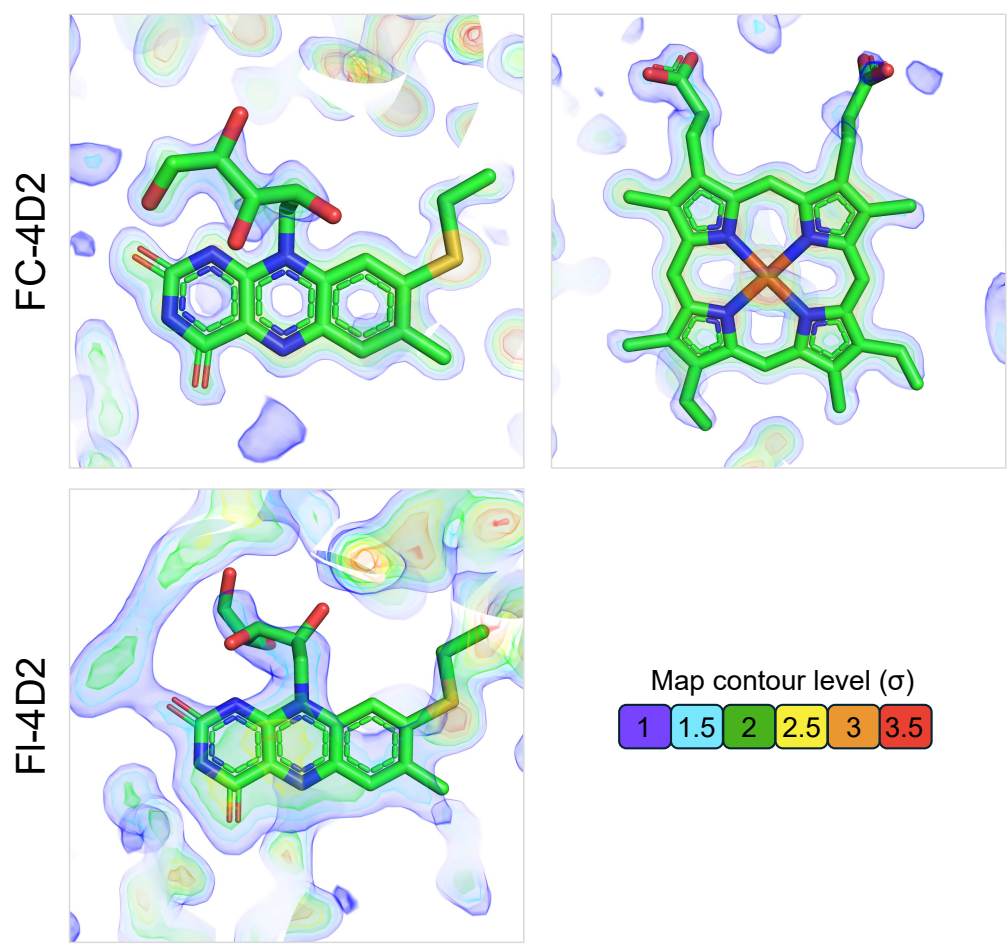

**Supplementary Figure S12:** $F_o-F_c$ OMIT maps of bound ligands in FC-4D2 and Fl-4D2 crystal structures. Contour $\sigma$ values are indicated by the scale at the bottom right of the images. Each ligand is homogenously bound, including the FC-4D2 heme despite previous 4D2-derived structures demonstrating multiplicity in heme plane ligation.<sup>2</sup>

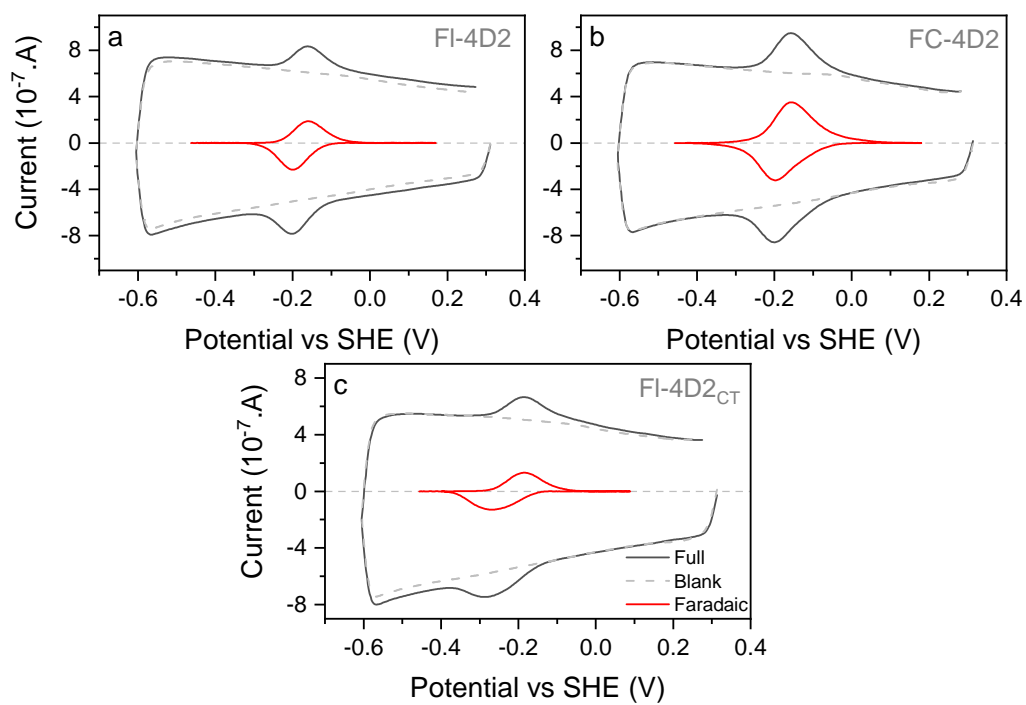

**Supplementary Figure S13:** PFE Cyclic voltammograms of (a) Fl-4D2, (b) FC-4D2 and (c) FI-4D2<sub>CT</sub> at 0.01 $\text{V.s}^{-1}$, pH 8.6. Faradaic currents (red) were subtracted from raw collected data ('Full', black) through baseline subtraction. CVs of blank electrodes (prepared with 20 mM CHES, 100 mM KCl, pH 8.6 buffer only) presented as grey dashed lines.

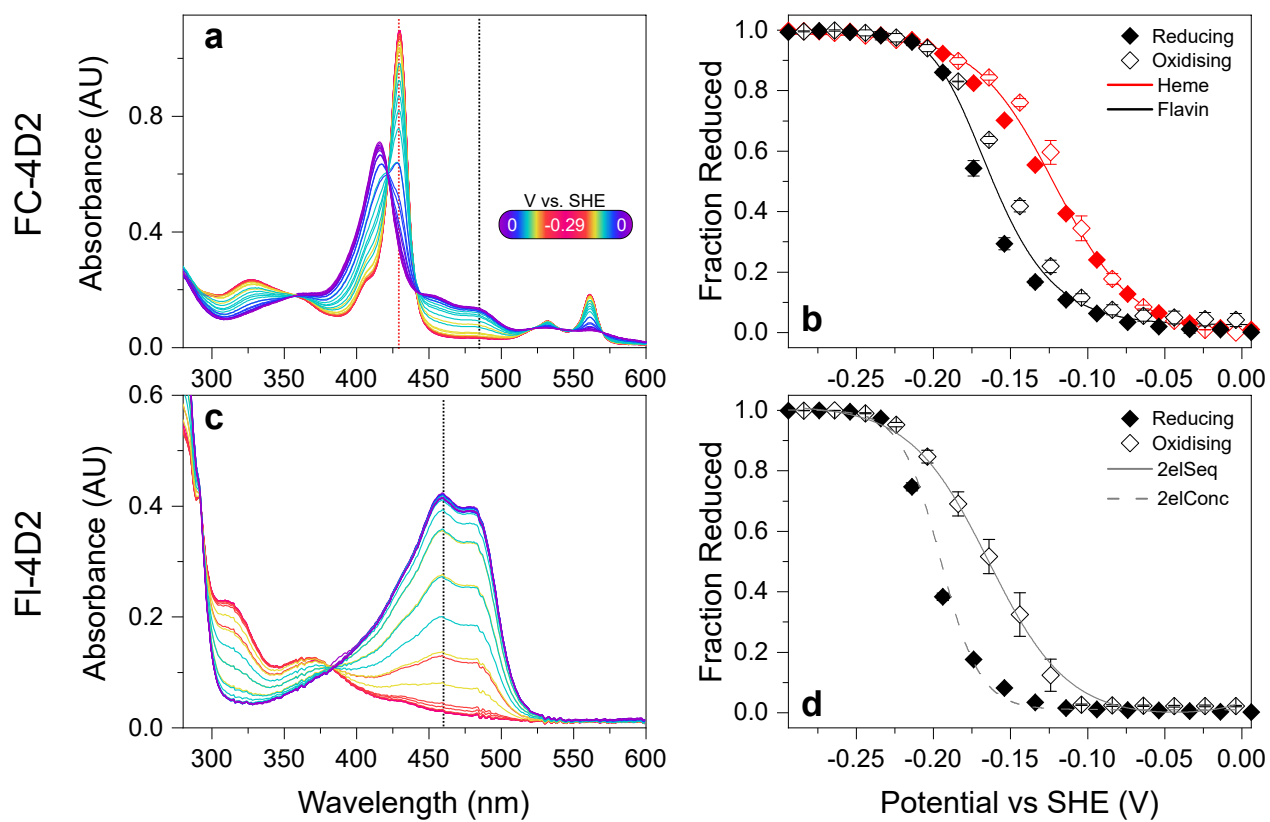

**Supplementary Figure S14:** OTTLE titrations of FC-4D2 and Fl-4D2 at pH 8.6, with equilibration times of 30 mins per 20 mV applied potential. (a) Absorbance of FC-4D2 as a function of applied potential (colourbar). Red and black dotted lines represent the wavelengths assessed for the heme and flavin titration respectively. (b) Potentiometric titrations of FC-4D2 heme (red) and flavin (black) presented as a fraction of reduced population and fit to a $1e^-$ (Eq. S13) and sequential $2 \times 1e^-$ (Eq. S14) Nernst equation respectively. Filled and open diamonds represent reducing and oxidising directions respectively. (c) Spectra as a function of applied potential of Fl-4D2, coloured identical to (a) with dotted line indicating wavelength assessed in (d). (d) Potentiometric titration of Fl-4D2 flavin, with reducing sweep (filled diamond) fit to a concerted $2e^-$ Nernst equation (2elConc, Eq. S15, dotted line) and oxidising (open diamond) sweep fit to a sequential $2 \times 1e^-$ Nernst equation (2elSeq, Eq. S14, solid line).

#### S1.2 FI-4D2<sub>CT</sub>

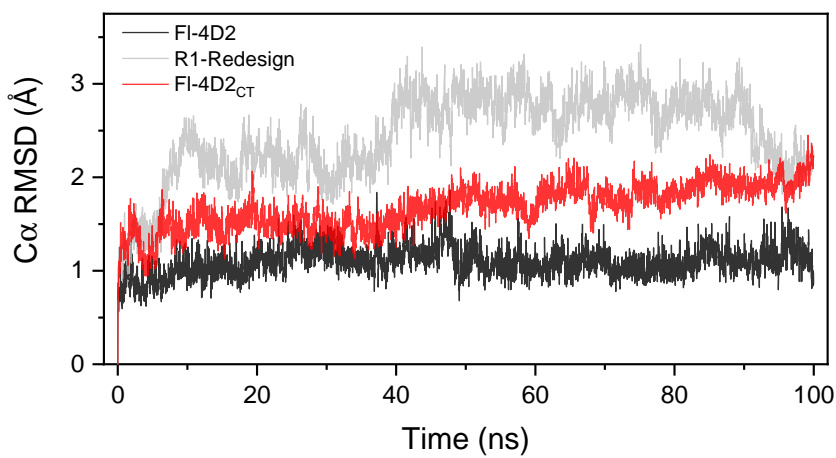

**Supplementary Figure S15:** Cα RMSD of Fl-4D2, R1-Redesign and FI-4D2<sub>CT</sub> across 100 ns MM MD.

| Electronic Transition | Avg EE (eV) | Avg $f_{osc}$ | Avg CT | | |
| --- | --- | --- | --- | --- | --- |
| **R1-Redesign** |  |  | 8-dMeRF to Phe27 | 8-dMeRF to Phe99 | Max |
| S<sub>1</sub> | 3.44 | 0.252 | 0.018 | 0.009 | 0.019 |
| S<sub>2</sub> | 3.62 | 0.192 | 0.034 | 0.009 | 0.034 |
| S<sub>3</sub> | 3.88 | 0.068 | 0.126 | 0.022 | 0.135 |
| S<sub>4</sub> | 4.11 | 0.063 | 0.134 | 0.062 | 0.179 |
| **FI-4D2<sub>CT</sub>** |  |  | 8-dMeRF to Trp27 | 8-dMeRF to Trp99 | Max |
| S<sub>1</sub> | 2.95 | 0.030 | 0.412 | 0.461 | 0.850 |
| S<sub>2</sub> | 3.30 | 0.068 | 0.319 | 0.389 | 0.628 |
| S<sub>3</sub> | 3.48 | 0.119 | 0.247 | 0.194 | 0.362 |
| S<sub>4</sub> | 3.67 | 0.086 | 0.250 | 0.183 | 0.361 |
| **FI-4D2** |  |  | 8-dMeRF to Gly27 | 8-dMeRF to Leu99 | Max |
| S<sub>1</sub> | 3.51 | 0.252 | 0.004 | 0.001 | 0.004 |
| S<sub>2</sub> | 3.73 | 0.273 | 0.005 | 0.002 | 0.005 |
| S<sub>3</sub> | 4.25 | 0.073 | 0.009 | 0.002 | 0.009 |
| S<sub>4</sub> | 4.47 | 0.050 | 0.008 | 0.002 | 0.008 |

**Supplementary Table S2:** Tables displaying calculated CT values between the 8-dMeRF isoalloxazine ring and aromatic residues (top) Phe27 and Phe99 for R1-Redesign, (middle) Trp27 and Trp99 in FI-4D2<sub>CT</sub> and (bottom) the corresponding residues, Gly27 and Leu99, in FI-4D2. CT values are between 0 and 1: 0 being no charge transfer between 8-dMeRF and specified residue(s) and 1 being complete charge-transfer. The excitation energies (EE), oscillator strengths ($f_{osc}$), and CT values are shown for each low-lying electronic transition. For each transition, the mean of these values were calculated over 100 snapshots.

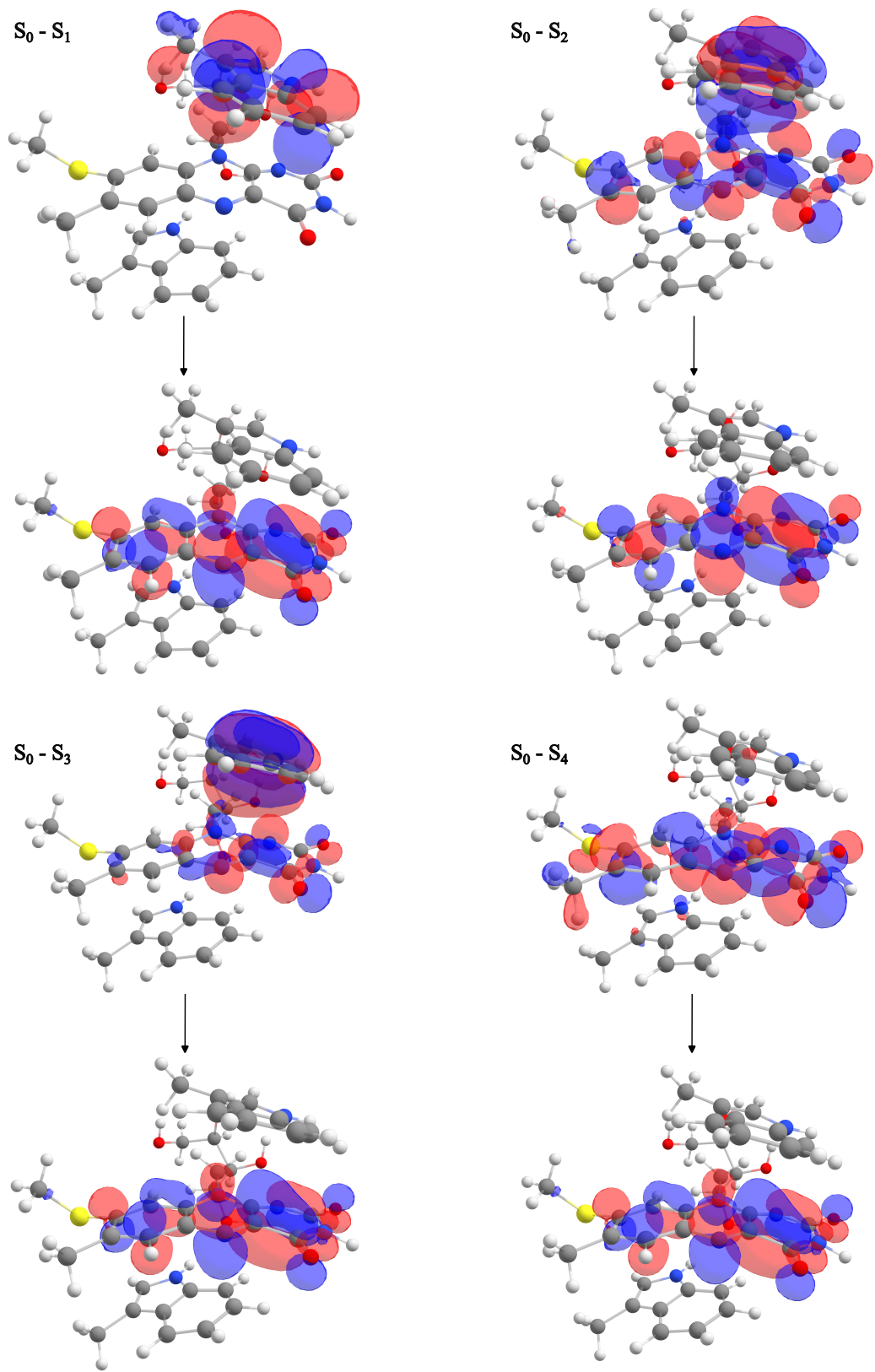

**Supplementary Figure S16:** NTOs from a sample QM/MM snapshot for low-lying transitions for FI-4D2<sub>CT</sub>. The QM region contains 8-dMeRF and the sandwiching Trp27 and Trp99 residues. The arrows connect the hole and particle NTOs associated with each electronic excitation.

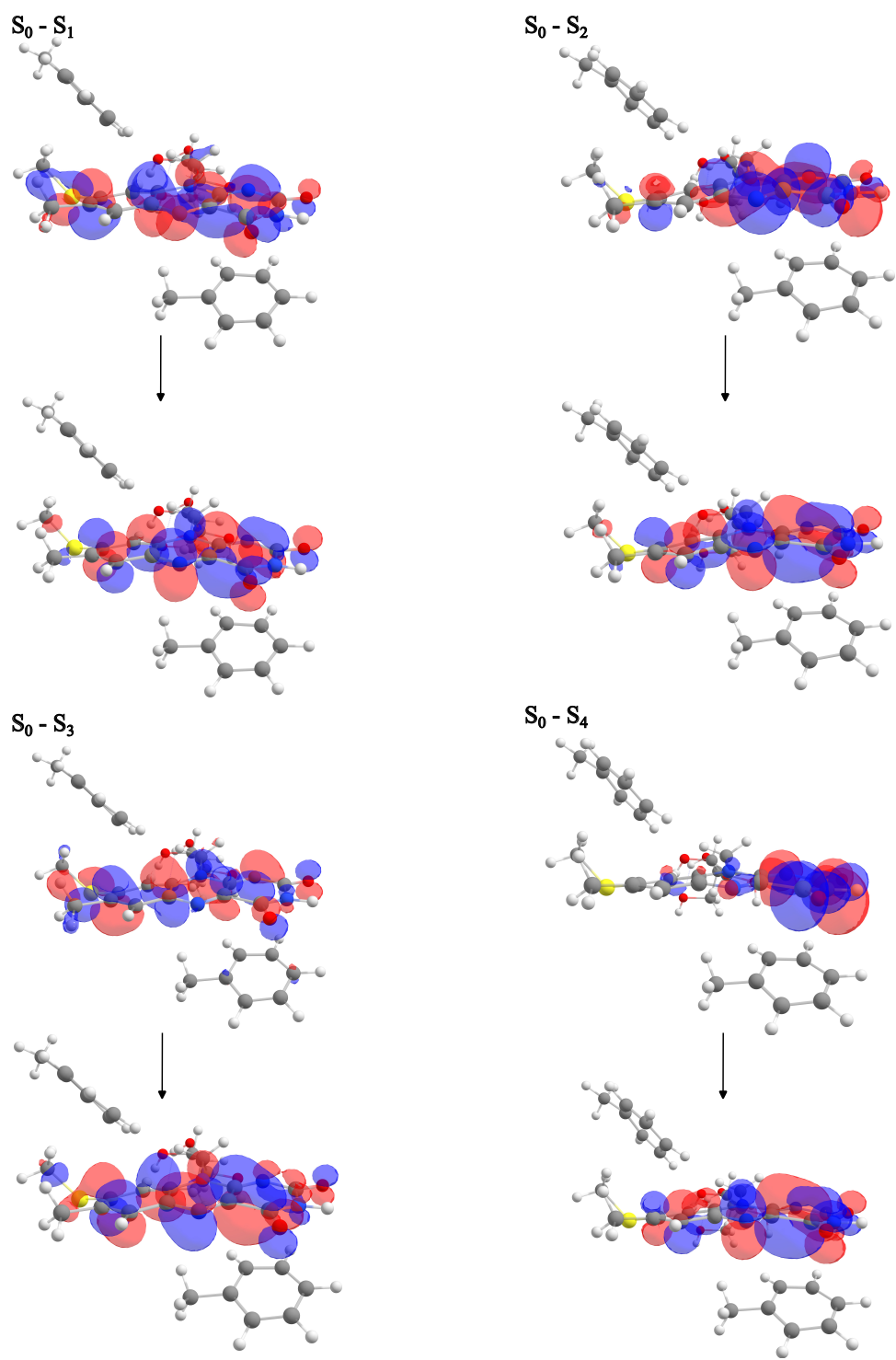

**Supplementary Figure S17:** NTOs from a sample QM/MM snapshot for low-lying transitions for R1-Redesign. The QM region contains 8-dMeRF and the sandwiching Phe27 and Phe99 residues. The arrows connect the hole and particle NTOs associated with each electronic excitation.

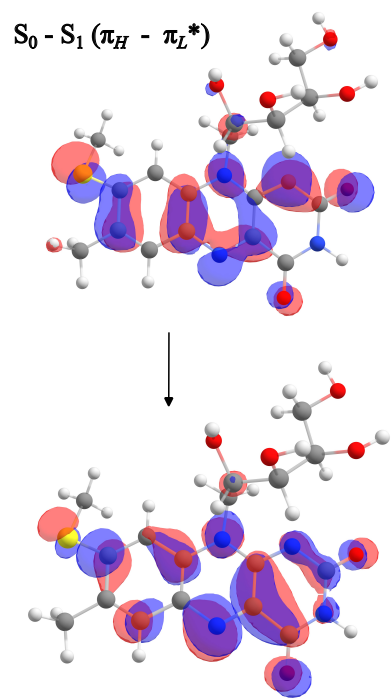

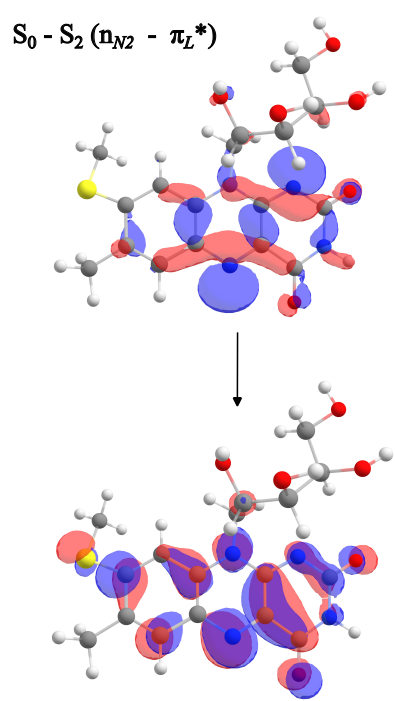

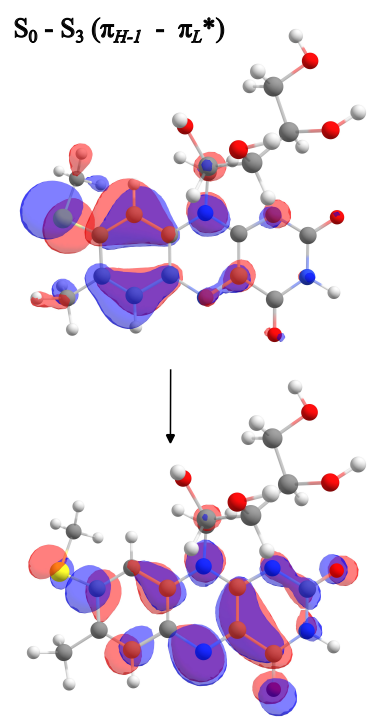

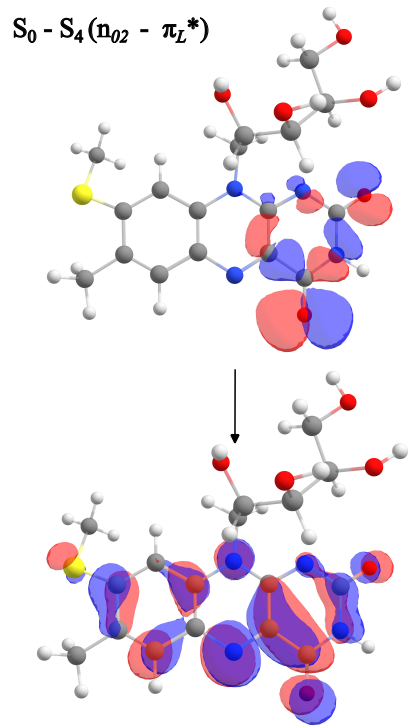

**Supplementary Figure S18:** NTOs from a sample QM/MM snapshot for low-lying transitions for Fl-4D2. The arrows connect the hole and particle NTOs associated with each electronic excitation.

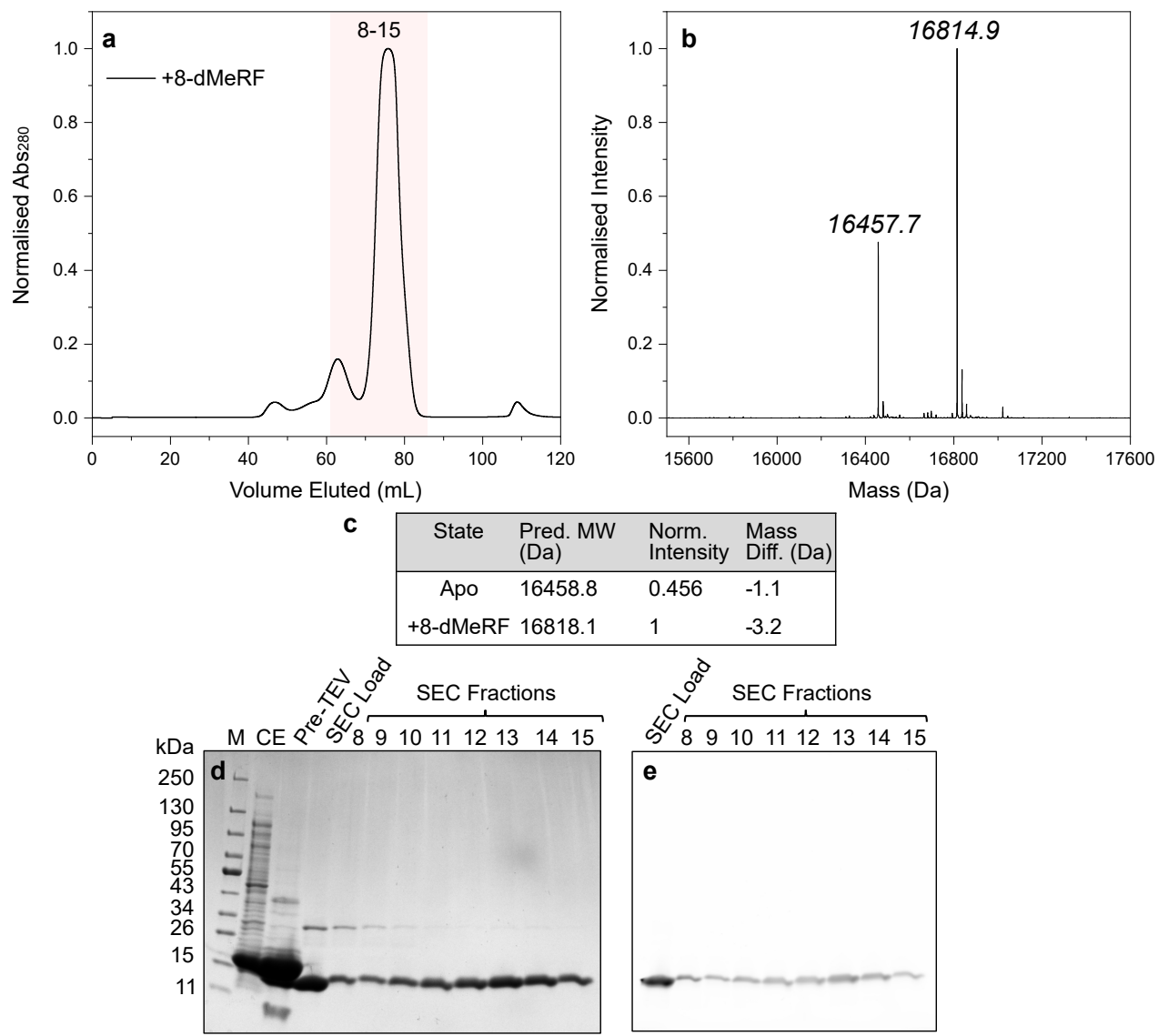

**Supplementary Figure S19:** Analytical characterisation of FI-4D2<sub>CT</sub>. (a) SEC chromatogram of FI-4D2<sub>CT</sub> in the holo state ('+8-dMeRF', black). Highlighted section represents range, fractions 8 through 15, sampled for analysis in SDS-PAGE. (b) Deconvoluted ESI-MS spectrum of holo FI-4D2<sub>CT</sub>, with the mass of most prominent species indicated. (c) Predicted mass of each FI-4D2<sub>CT</sub> state, with normalised intensity and mass difference relative to the corresponding ESI-MS peak. (d) SDS-PAGE of FI-4D2<sub>CT</sub> throughout purification, with the corresponding fluorescent image of the same gel upon excitation with 473 nm laser (e).

#### S2 Photoreduction

##### S2.1 Continuous-wave photoreduction

#### S2.1.1 FI-4D2

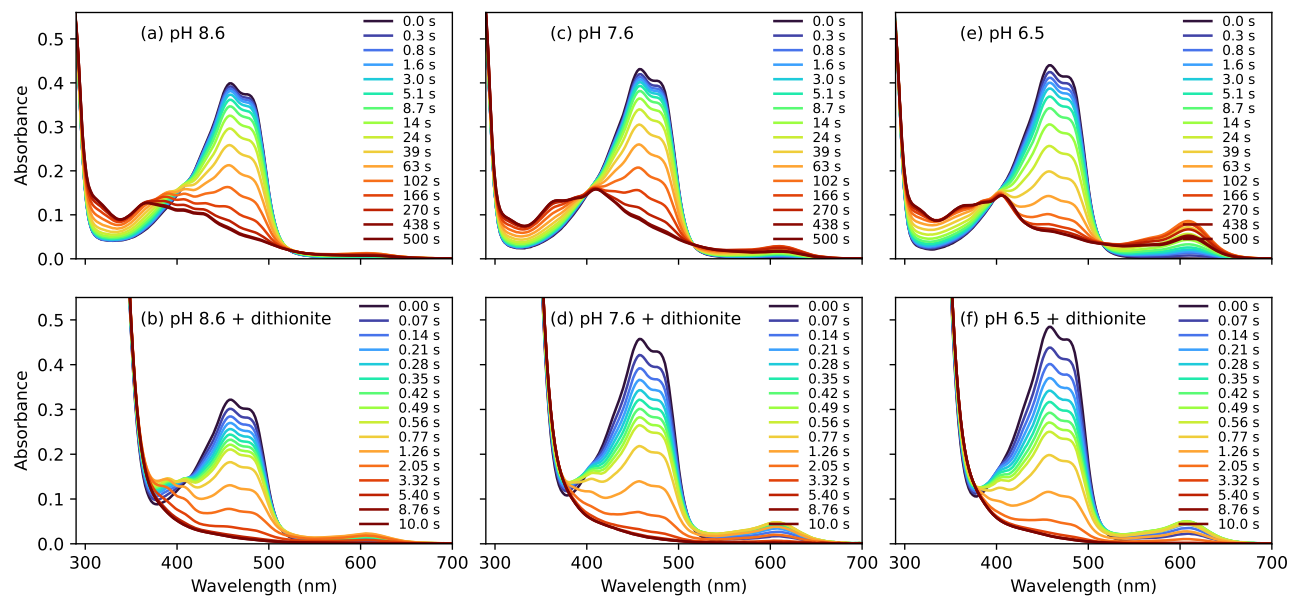

**Supplementary Figure S20:** Steady-state absorbance of Fl-4D2 after different cw irradiation times, in buffers CHES (a,b), HEPES (c,d), and MES (e,f), and with dithionite (b,d,f).

Photoreduction data was collected under cw white-light irradiation for Fl-4D2 in three different buffers: CHES (pH 8.6), HEPES (pH 7.6), and MES (pH 6.5). A small amount of oxidized flavin (8-dMeRF) remains even after 500 s of irradiation in these experiments, so data were also collected with the addition of the strong reducing agent dithionite (which absorbs <400 nm)<sup>3</sup>, to obtain the spectrum of the fully-reduced species.

With irradiation, 8-dMeRF (F) is photoreduced first to semiquinone form, then to hydroquinone form. Two protonation states exist for each form: $\text{FH}^\bullet$ and $\text{F}^{\bullet-}$ in the semiquinone form, $\text{FH}^-$ and $\text{FH}_2$ in the hydroquinone form (Fig.3a). The visible absorption spectra of the various flavoproteins at any irradiation time can thus be fit to a linear combination of the absorption of each of the redox states, plus dithionite if it is included. The basis spectra of each redox state were extracted as follows:

• The absorption spectrum of 8-dMeRF ($\epsilon_F(\lambda)$) was extracted from the steady-state Fl-4D2 absorption spectrum without dithionite at 0 s of irradiation:

$$\epsilon_F(\lambda) = A_{\text{FI-4D2}}(0 \text{ s}, \lambda) / c_F(0 \text{ s})l \quad (\text{S1})$$

The concentration of 8-dMeRF in the protein was not known precisely, so the concentration $c_F(0 \text{ s})$ was scaled to match the peak of $\epsilon_F(\lambda)$ to the literature value for riboflavin in solution ($\sim 12300 \text{ M}^{-1}\text{cm}^{-1}$).<sup>4</sup>

• The dithionite spectrum ($\epsilon_D(\lambda)$) was extracted from the Fl-4D2 with dithionite spectrum at 0 s, subtracting away an appropriate amount of riboflavin.
• The hydroquinone spectrum ($\epsilon_{(\text{FH}^-/\text{FH}_2)}(\lambda)$) was extracted from Fl-4D2 with dithionite spectrum at 10 s, by which time the spectral evolution was complete and no species associated with oxidised or semiquinone flavin are observed, indicating only the hydroquinone form of 8-dMeRF and dithionite were present. The spectral shape at 10 s was the same regardless of buffer, which could either indicate that the same relative

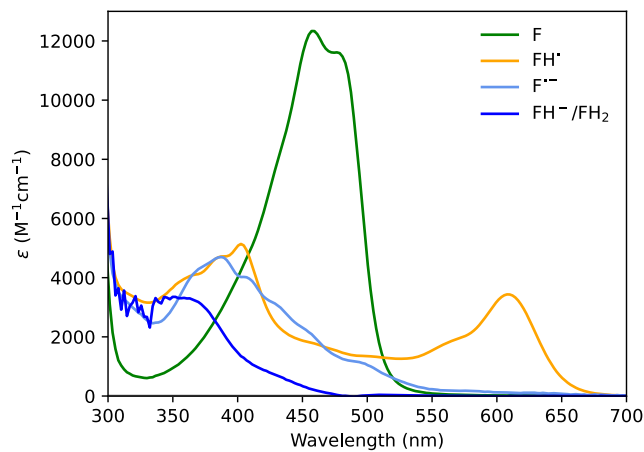

**Supplementary Figure S21:** Basis spectra for 8-dMeRF (F), 8-dMeRF semiquinones (F<sup>•-</sup> and FH<sup>•</sup>) and 8-dMeRF hydroquinone (FH<sup>-</sup>/FH<sub>2</sub>).

amounts of FH<sup>-</sup> and FH<sub>2</sub> are formed in each buffer with dithionite present, or that the absorption of the two protonation states are too similar to distinguish in the UV-visible region. We hence do not attempt to identify separate basis spectra for FH<sup>-</sup> and FH<sub>2</sub>, and treat them as one species. The similarity spectra of FH<sup>-</sup> and FH<sub>2</sub> is consistent with the literature.<sup>5</sup>

• The semiquinone spectra $\epsilon_{(FH^{\bullet})}(\lambda)$ and $\epsilon_{(F^{\bullet-})}(\lambda)$ were extracted from the samples with no dithionite at early irradiation times (88 s), before any significant quantities of hydroquinone had formed. The spectral profiles around these times varied with pH, likely due to varying proportions of the protonated (FH<sup>•</sup>) and non-protonated (F<sup>•-</sup>) semiquinone. Since the pKa of semiquinone protonation is approximately 8,<sup>6-8</sup> we assign the semiquinone spectrum extracted from pH 6.5 as predominantly FH<sup>•</sup>, and the spectrum extracted from pH 8.6 as predominantly F<sup>•-</sup>. Note that the spectra are likely not pure: i.e. $\epsilon_{(FH^{\bullet})}(\lambda)$ likely contains contributions from F<sup>•-</sup> and vice versa, but they are nonetheless useful for indicating differences in the proportions of FH<sup>•</sup> and F<sup>•-</sup> present.

The extracted basis spectra are shown in Fig. S21. They agree well with previously reported spectra for FMN, FAD and riboflavin.<sup>4,5,9</sup> The spectrum of FI-4D2 in each buffer at each irradiation time could thus be fit as a linear combination of these five basis spectra:

$$A(\lambda, t) = \epsilon_F(\lambda)c_F(t) + \epsilon_{(FH^{\bullet})}(\lambda)c_{(FH^{\bullet})}(t) + \epsilon_{(F^{\bullet-})}(\lambda)c_{(F^{\bullet-})}(t) + \epsilon_{(FH^-/FH_2)}(\lambda)c_{(FH^-/FH_2)}(t) + \epsilon_D(\lambda)c_D(t) \quad (S2)$$

Where $c_x(t)$ is the concentration of species $x$ after irradiation time $t$. The resulting concentrations are shown in Figs S22 and S23, and the fits of the linear combination are in Figs S24 and S25. Note that the presence of dithionite likely decreases the pH, so the HEPES buffer solution (“pH 7.6”) may actually be significantly lower than the semiquinone protonation pKa (~7) with dithionite, so only FH<sup>•</sup> is needed to fit the spectra. Without dithionite, the HEPES pH is close enough to the pKa that both F<sup>•-</sup> and FH<sup>•</sup> are needed to reproduce the data. Without dithionite, we observe the greatest amount of unprotonated semiquinone (F<sup>•-</sup>) at the highest pH (8.6, CHES) as expected, and similarly the greatest amount of FH<sup>•</sup> at the lowest pH (6.5, MES), whilst the mid-range pH (7.6, HEPES), contains both.

There are some small discrepancies between fits of the deconvoluted basis spectra and the experimental data, particularly at long irradiation times for samples without dithionite. This may partly arise from (1) the basis spectra of F<sup>•-</sup> and FH<sup>•</sup> are not entirely “pure”, so the exact proportions present cannot be reproduced, (2) different proportions of FH<sup>-</sup>/FH<sub>2</sub> present than in the dithionite samples (where the FH<sup>-</sup>/FH<sub>2</sub> spectra were extracted), or (3) competing photodegradation pathways, such as the formation of lumiflavin or lumichrome (with a methyl group or hydrogen replacing the ribityl tail, respectively).<sup>10,11</sup>

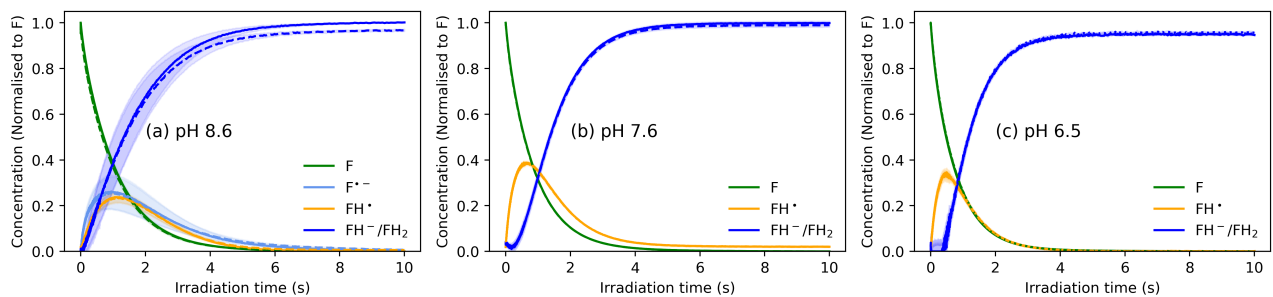

**Supplementary Figure S22:** Deconvoluted concentrations of 8-dMeRF (F), 8-dMeRF semiquinones ($F^{\bullet-}$ and $FH^{\bullet}$) and 8-dMeRF hydroquinone ($FH^-/FH_2$) as a function of irradiation time under a cw light source, for Fl-4D2 in a CHES (a), HEPES (b) or MES (c) buffers, **with dithionite**. Repeated experiments are shown in dashed and/or dotted lines. Shaded areas are 95% confidence interval of the fitted concentrations.

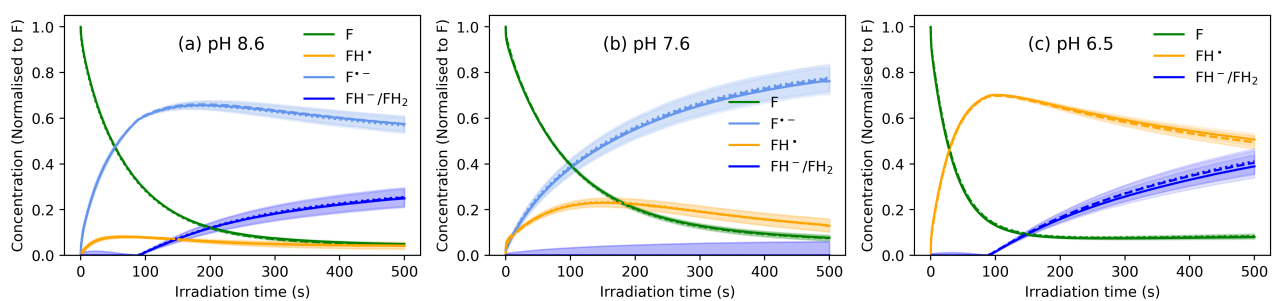

**Supplementary Figure S23:** Deconvoluted concentrations of 8-dMeRF (F), 8-dMeRF semiquinones ($F^{\bullet-}$ and $FH^{\bullet}$) and 8-dMeRF hydroquinone ($FH^-/FH_2$) as a function of irradiation time under a cw lamp, for Fl-4D2 in a CHES (a), HEPES (b) or MES (c) buffer, **without dithionite**. Shaded areas are 95% confidence interval of the fitted concentrations.

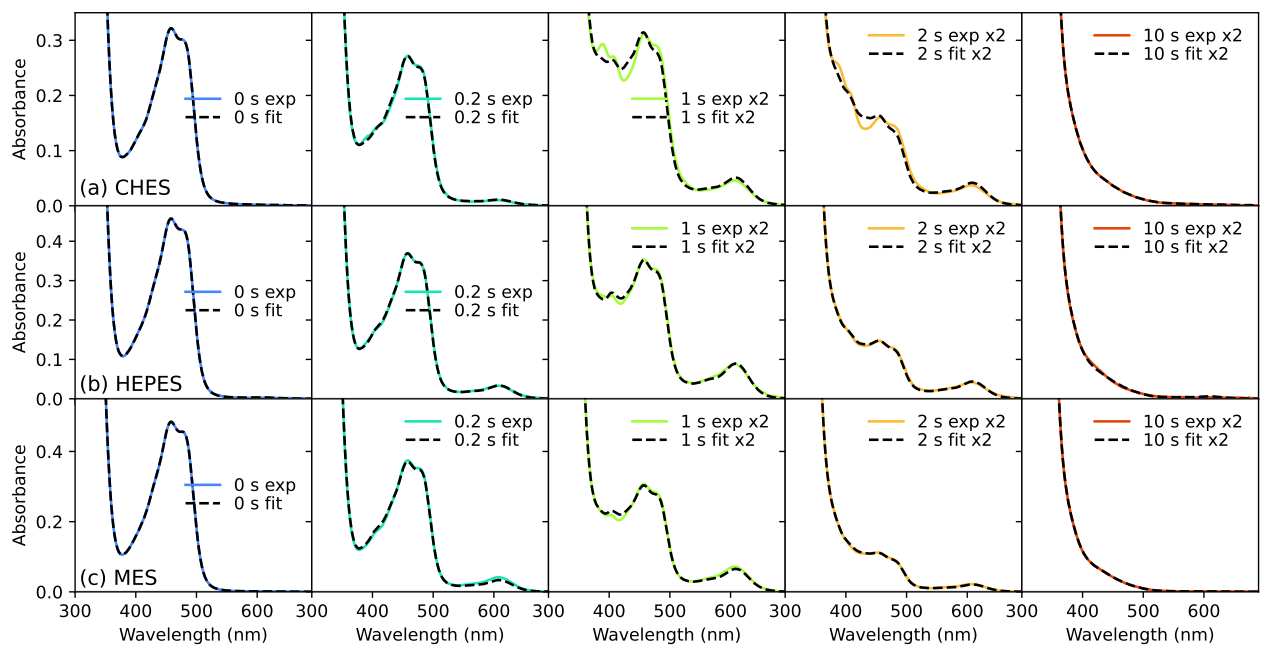

**Supplementary Figure S24:** Fits of F, $F^{\bullet-}$, $FH^{\bullet}$, and $FH^-/FH_2$ basis spectra to the Fl-4D2 **with dithionite** in a CHES (a), HEPES (b) or MES (c) buffer, at selected irradiation times.

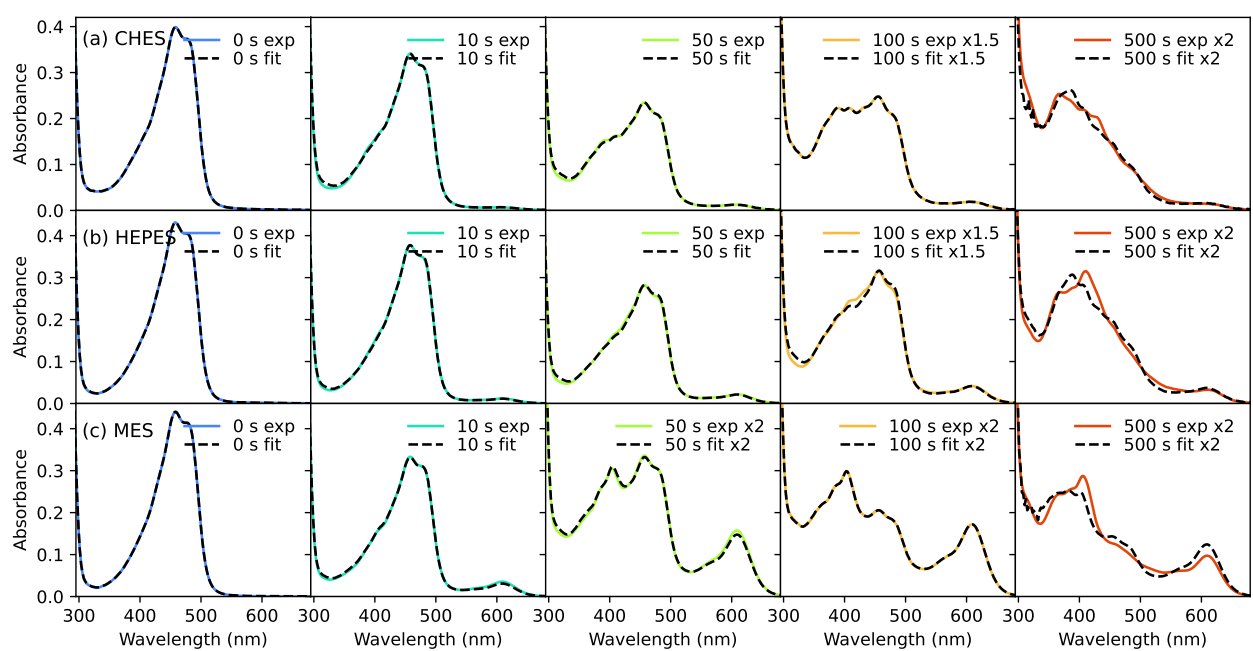

**Supplementary Figure S25:** Fits of F, $F^{\bullet-}$, $FH^{\bullet}$, and $FH^-/FH_2$ basis spectra to the Fl-4D2 **without dithionite** in a CHES (a), HEPES (b) or MES (c) buffer, at selected irradiation times.

#### S2.1.2 FI-4D2<sub>CT</sub>

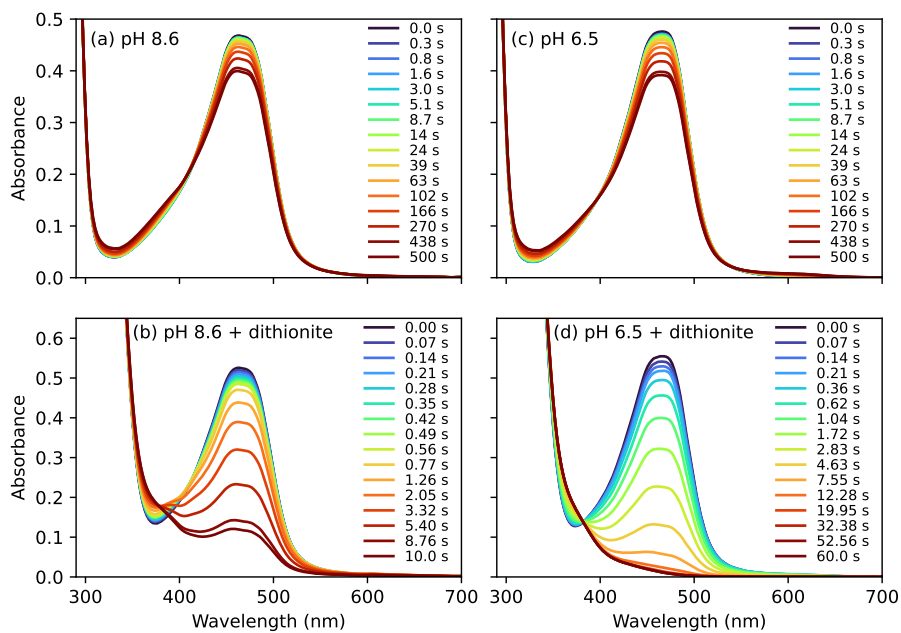

**Supplementary Figure S26:** Steady-state absorbance of FI-4D2<sub>CT</sub> after different cw irradiation times, in buffers CHES (a,b), and MES (c,d), and with dithionite (b,d).

Since the 8-dMeRF absorption spectrum is different in FI-4D2<sub>CT</sub>, the basis spectrum of (oxidised) 8-dMeRF was redefined to fit the FI-4D2<sub>CT</sub> photoreduction, again using the spectrum after 0 s irradiation with no dithionite present. The hydroquinone spectrum was also extracted from samples with dithionite at 60 s irradiation time. Whilst it is likely that the FI-4D2<sub>CT</sub> semiquinone basis spectra also vary from Fl-4D2, since the flavin in the FI-4D2<sub>CT</sub> design undergoes little photoreduction without dithionite, it could not be isolated from the cw lamp irradiated data. Hence the semiquinone basis spectra extracted from Fl-4D2 were used to fit the data. Note that since only very small amounts of reduced flavin are formed in the absence of dithionite, the resolution of the photoproducts is low, and it is also possible that FI-4D2<sub>CT</sub> is degrading via some other pathway (e.g. ~lumichrome formation), rather than being photoreduced to semiquinone and hydroquinone forms.<sup>10,11</sup>

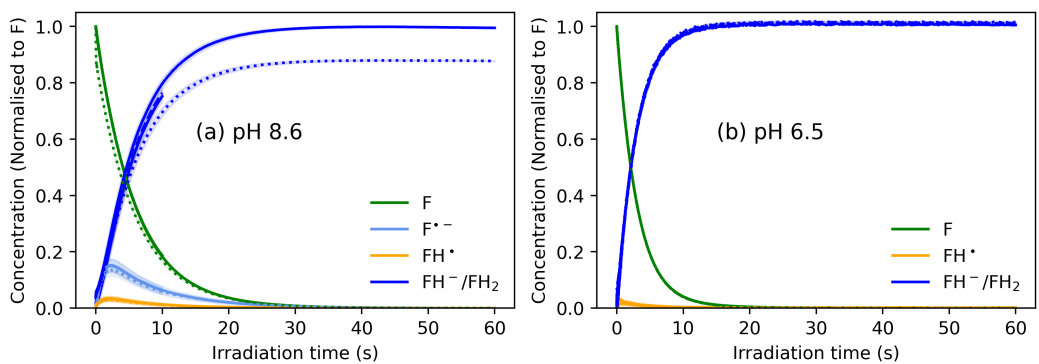

**Supplementary Figure S27:** Deconvoluted concentrations of 8-dMeRF (F), 8-dMeRF semiquinones (F<sup>•-</sup> and FH<sup>•</sup>) and 8-dMeRF hydroquinone (FH<sup>-</sup>/FH<sub>2</sub>) as a function of cw irradiation time, for FI-4D2<sub>CT</sub> in a CHES (a) or MES (b) buffer, **with dithionite**. Shaded areas are 95% confidence interval of the fitted concentrations.

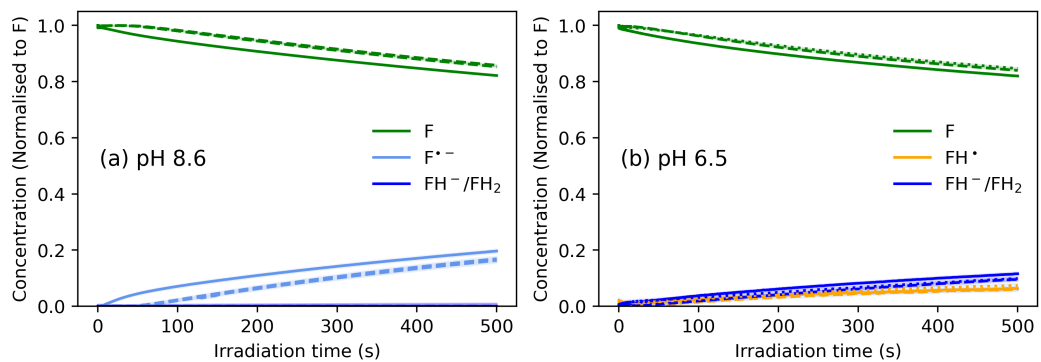

**Supplementary Figure S28:** Deconvoluted concentrations of 8-dMeRF (F), 8-dMeRF semiquinones ($F^{\bullet-}$ and $FH^{\bullet}$) and 8-dMeRF hydroquinone ($FH^-/FH_2$) as a function of cw irradiation time, for FI-4D2<sub>CT</sub> in a CHES (a) or MES (b) buffer, **without dithionite**. Shaded areas are 95% confidence interval of the fitted concentrations.

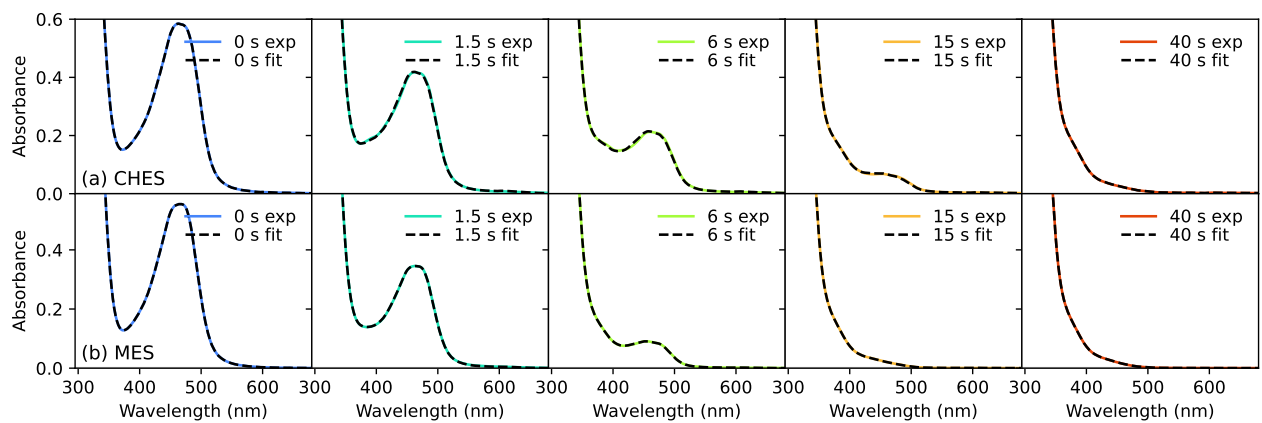

**Supplementary Figure S29:** Fits of F, $F^{\bullet-}$, $FH^{\bullet}$, and $FH^-/FH_2$ basis spectra to the FI-4D2<sub>CT</sub> **with dithionite** in a CHES (a) or MES (b) buffer, at selected irradiation times.

**Supplementary Figure S30:** Fits of F, $F^{\bullet-}$, $FH^{\bullet}$, and $FH^-/FH_2$ basis spectra to the FI-4D2<sub>CT</sub> **without dithionite** in a CHES (a) or MES (b) buffer, at selected irradiation times.

#### S2.2 10 kHz pulsed photoreduction

Both Fl-4D2 and FC-4D2 were also photoreduced with a 10 kHz pulsed laser at 480 nm (~200 fs FWHM, 30 mW). At 10 kHz there is 100 μs between pulses, over which at least 95% of excited states would have recovered to the ground state, according to TA data. Additionally, most of the flavin photoproducts absorb minimally at 480 nm. Hence, in contrast to the white-light cw irradiation, the pulsed 480 nm source will predominantly excite ground-state flavin. We also performed the pulsed photoreduction with the addition of a cw 633 nm light source (HeNe, 7 mW) to also excite the FH$^{\bullet}$ species or other photoproducts, to test if this affects the rate of hydroquinone formation. In both cases the flavin was still fully reduced to the hydroquinone, as observed for the cw irradiation. Further, the rate of photodegradation of FI-4D2 was unchanged with the addition of the 633 nm source, hence we can conclude the full photoreduction is not reliant on the double excitation of flavin excited states or other photoproducts.

The absorption spectra after pulsed irradiation were fit with the same basis spectra as used for the cw data (Fig. S21). Interestingly, the red-absorbing FH$^{\bullet}$ was not observed in the pulsed photoreduction experiments. There was some delay (~60 s) between switching the irradiation off and collecting the absorption spectrum for the pulsed photoreduction experiments, whereas the cw irradiation measurements were collected in situ. Hence it is possible FH$^{\bullet}$ was depleted before it could be detected for pulsed photoreduction experiments.

We note that whilst the additional 633 nm irradiation had no effect on Fl-4D2, it did drive the heme reduction in FC-4D2 further to completion. This suggests there may be an additional photoinduced electron transfer pathway from the excited state of FH$^{\bullet}$ to heme.

### S2.2.1 FI-4D2

**Supplementary Figure S31:** Steady-state absorbance of Fl-4D2 at different irradiation times under 480 nm 10 kHz pulsed irradiation (a), and 480 nm pulsed plus 633 nm continuous wave irradiation (b), for CHES buffer (pH 8.6) and without dithionite.

**Supplementary Figure S32:** Deconvoluted concentrations of 8-dMeRF ($F$), 8-dMeRF semiquinone ($F^{\bullet-}$), 8-dMeRF hydroquinone ($FH^-/FH_2$) as a function of irradiation time for Fl-4D2 in a CHES buffer without dithionite, under (a) 480 nm 10 kHz pulsed irradiation, and (b) 480 nm pulsed plus 633 nm cw irradiation. Shaded area indicates a 95% confidence interval. (c) shows the concentrations on the same axes, with 480 nm irradiation in circles and full lines, and 480 nm + 633 nm irradiation in triangles and dashed lines.

**Supplementary Figure S33:** Fits of $F$, $F^{\bullet-}$, $FH_2$, and $FH^-/FH_2$ (same as though extracted for the cw spectra) to the Fl-4D2 without dithionite in a CHES buffer, at selected 480 nm 10 kHz pulsed irradiation times (a), and 480 nm pulsed plus 633 nm cw irradiation times (b).

## S2.2.2 FC-4D2

**Supplementary Figure S34:** Steady-state absorbance of FC-4D2 at different irradiation times under 480 nm 10 kHz pulsed irradiation (a), and 480 nm pulsed plus 633 nm cw irradiation (b), for CHES buffer (pH 8.6) and without dithionite.

**Supplementary Figure S35:** Deconvoluted concentrations of 8-dMeRF (F), 8-dMeRF semiquinone (F<sup>•-</sup>), 8-dMeRF hydroquinone (FH<sup>-</sup>/FH<sub>2</sub>), and oxidized and reduced heme, as a function of irradiation time for FC-4D2 in a CHES buffer (pH 8.6) without dithionite, under 480 nm 10 kHz pulsed irradiation (a), and 480 nm pulsed plus 633 nm cw irradiation (b). Shaded area indicates a 95% confidence interval. (c) and (d) show the concentrations on the same axes, with 480 nm irradiation in circles and full lines, and 480 nm + 633 nm irradiation in triangles and dashed lines.

**Supplementary Figure S36:** Fits of F, F<sup>•-</sup>, FH<sup>-</sup>/FH<sub>2</sub> and oxidised and reduced heme (same as those extracted for the cw spectra) to the FC-4D2 without dithionite in a CHES buffer, at selected 480 nm 10 kHz pulsed irradiation times (a), and 480 nm pulsed plus 633 nm continuous wave irradiation times (b).

#### S2.3 Excitation rate calculations

To determine the efficiency of the photoreduction of flavin in Fl-4D2, the excitation rate was calculated from the excitation fluence and the initial ground state absorption.

The power measured on a power meter with a 9.5 mm diameter (70.88 mm<sup>2</sup> area) was 30 mW (0.03 J/s). The density of photons incident on the sample can thus be calculated as:

$$\begin{aligned} \text{power density} &= \frac{0.03 \text{ Js}^{-1}}{70.88 \text{ mm}^2} \\ &= 4.23 \times 10^{-4} \text{ J s}^{-1} \text{ mm}^{-2} \end{aligned} \quad (\text{S3})$$

$$\begin{aligned} \text{energy per photon} &= \frac{hc}{480 \text{ nm}} \\ &= 4.14 \times 10^{-19} \text{ J} \end{aligned} \quad (\text{S4})$$

$$\begin{aligned} \text{incident photon density} &= \frac{\text{power density}}{\text{energy per photon}} \\ &= 1.022 \times 10^{15} \text{ s}^{-1} \text{ mm}^{-2} \end{aligned} \quad (\text{S5})$$

The initial absorbance of the sample was 0.126 in a 1 mm pathlength cuvette. Hence, the density of photons absorbed can be calculated as:

$$\begin{aligned} \text{absorbed photon density} &= \text{incident photon density} \times (1 - 10^{-\text{absorbance}}) \\ &= 2.574 \times 10^{14} \text{ s}^{-1} \text{ mm}^{-2} \end{aligned} \quad (\text{S6})$$

then,

$$\begin{aligned} \text{exciton density} &= \text{absorbed photon density} \\ &= 2.574 \times 10^{14} \text{ s}^{-1} \text{ mm}^{-2} \\ &= 4.27 \times 10^{-4} \text{ molL}^{-1} \text{ s}^{-1} \\ &= k_{\text{ex}}[\text{S}_0](t = 0) \end{aligned} \quad (\text{S7})$$

Hence, for an initial concentration ($[S_0](t = 0)$) of $1.08 \times 10^{-5} \text{ molL}^{-1}$,

$$k_{\text{ex}} = 39.7 \text{ s}^{-1} \quad (\text{S8})$$

Given 88.55% of the Flavin ground states is reduced in 100 s of 10 kHz 30 mW irradiation, we can approximate the amount excited states reduced per pulse. Following from calculations above:

$$\begin{aligned} \text{excitation density} &= 4.27 \times 10^{-4} \text{ molL}^{-1} \text{s}^{-1} \\ \text{rep rate} &= 10000 \text{ s}^{-1} \\ \therefore \text{excitations per pulse} &= 4.27 \times 10^{-8} \text{ molL}^{-1} \end{aligned} \quad (\text{S9})$$

This corresponds to 0.4% of the initial $S_0$ concentration ($1.08 \times 10^{-5} \text{ molL}^{-1}$). In 100 s, there are $10^6$ pulses, and 89% of the sample is reduced i.e.

$$\begin{aligned} \text{molecules reduced in 100 s} &= 9.54 \times 10^{-6} \text{ molL}^{-1} \\ \therefore \text{molecules reduced per pulse} &= \frac{9.54 \times 10^{-6} \text{ molL}^{-1}}{10^6} \\ &= 9.54 \times 10^{-12} \text{ molL}^{-1} \end{aligned} \quad (\text{S10})$$

$$\begin{aligned} \text{percent of excited states reduced per pulse} &= \frac{\text{molecules reduced per pulse}}{\text{excitons per pulse}} \times 100 \\ &= 0.022\% \end{aligned} \quad (\text{S11})$$

Note this calculation represents the number of excitons reduced per pulse *on average* over 100 s of irradiation. In reality less excitons are excited over time as the sample is photoreduced, so 0.022% is an lower estimate of the percentage at 0 s, and upper estimate of the percentage at 100 s.

##### S3 Transient Absorption Spectroscopy

**Comparison of FI-4D2 and FI-4D2<sub>CT</sub> lifetimes** Figure S37 shows the transient absorption (TA) spectra of FI-4D2 and FI-4D2<sub>CT</sub> normalized to the same amplitude at 1 ps. By 50 ps, there is significantly more signal for Fl-4D2 than FI-4D2<sub>CT</sub>. By 100 ps, negligible signal remains for FI-4D2<sub>CT</sub>, whilst 6% of the signal is still present for FI-4D2.

**Spectral profile of FI-4D2<sub>CT</sub>** As Fig. S37 shows, the TA spectral profile of FI-4D2<sub>CT</sub> is different to Fl-4D2, even at very early times. In particular, we note the lack of any stimulated emission features from the flavin singlet excited state ($^1F^*$), and an additional broad positive photoinduced absorption (PIA) between 500–700 nm. It has been well documented in the literature that tryptophan radical cations absorb in this region, as shown in Fig. S38.<sup>12,13</sup> Thus, based on this PIA and QM/MM predictions we assign the product of excitation as a CT state between between the flavin and tryptophan residues.

Other TA studies of flavoproteins have observed the formation of flavin:tryptophan radical pairs $F^{\bullet-} \text{TrpH}^{\bullet+}$ (the product of electron transfer) or $FH^{\bullet} \text{Trp}^{\bullet}$ (the product of electron and proton transfer) that appear as an exact linear combination of steady-state flavin semiquinone and tryptophan cation spectra.<sup>12</sup> However, as Fig. S38 shows, the FI-4D2<sub>CT</sub> spectrum is not an exact match for either radical pair. The PIA from 650–700 nm cannot be accounted for by the steady-state spectrum of either $\text{TrpH}^+$ or $FH^{\bullet}$, and the sharp PIA centred at 390 nm is much more consistent with the $^1F^*$ absorption observed in Fl-4D2. We thus consider the product of exciting FI-4D2<sub>CT</sub> to be a singlet CT state, rather than a well-separated radical pair. Since no spectral evolution is observed, this CT state either forms extremely rapidly from $^1F^*$ within the 200 fs instrument response function, or else is the direct product of excitation (i.e. FI-4D2<sub>CT</sub> is excited directly to the CT state).

**Supplementary Figure S37:** Comparison of the amplitudes of the TA spectrum of FI-4D2 and FI-4D2<sub>CT</sub> at 1 ps, 50 ps, and 100 ps, showing Fl-4D2 has a long-lived population that is not present in FI-4D2<sub>CT</sub>.

**Supplementary Figure S38:** Transient absorption spectrum of FI-4D2<sub>CT</sub> at 1 ps, compared with the steady-state spectra of two potential radical pairs: (a) TrpH<sup>•+</sup> and F<sup>•-</sup>, or (b) Trp<sup>•</sup> and FH<sup>•</sup>. The flavin semiquinone spectra F<sup>•-</sup> and FH<sup>•</sup> are taken from photoreduction experiments of FI-4D2, and Tryptophan spectra are taken from Gravell *et al.*<sup>12</sup>

**Deconvolution of FI-4D2<sub>CT</sub> and comparison with solution** Unlike FI-4D2<sub>CT</sub>, the TA spectrum of FI-4D2 shows some spectral evolution over time. We thus performed a spectral deconvolution of both Fl-4D2 and riboflavin in DMSO solution, the latter having being reported previously,<sup>14</sup> to clarify the photophysics. Only two spectral components are necessary to describe the photophysics of riboflavin in solution: a singlet basis spectrum ($^1F^*$) extracted from early times (1 ps), and a triplet basis spectrum ($^3F^*$) extracted from late times (100 ns) (Fig. S39b). Fitting a linear combination of these basis spectrum to the full TA spectra at each delay time lead to the concentration profiles in dashed lines in Fig. S39d, demonstrating intersystem crossing of riboflavin in solution.<sup>14</sup>

The spectrum of FI-4D2 is significantly more complicated: at least four spectral components are necessary to fit the data, likely due to multiple conformations leading to multiple spectrally distinct populations. To enable a simple comparison with FI-4D2<sub>CT</sub> and riboflavin in solution, we performed a rough spectral deconvolution of this sample. We tentatively assign two protein populations: one which is susceptible to some rapid deactivation of the singlet excited state (A), and one that is not (B) (Fig. S40). We extract an excited singlet basis spectrum for the longer-lived population B, $^3F^*_B$, at 20 ps, after a the initial early-time spectral evolution was complete (Fig. S39a). The triplet basis spectrum for population B, $^1F^*_B$, was extracted at late times at which no stimulated emission from singlet states could be observed (40 ns). Two additional singlet basis spectra were necessary to fit the data before 20 ps, which we assign as singlet excited state and a vibrationally hot singlet excited state on population A, $^1F^*_A$ and $^1F^{**}_A$, respectively. For simplicity, we group these two excited states together in the kinetics in Fig. S39c. These assignments are supported by the stimulated emission features in the $^1F^*_A$ and $^1F^*_B$ spectra, and the similar shapes of the $^1F^*_B$ and $^3F^*_B$ spectra to that of riboflavin in DMSO.

The deconvoluted profiles in Fig. S39c shows that $^1F^*_A$ accounts for approximately 90% of the FI-4D2 excited state population, and decays much faster than FI-4D2<sub>CT</sub>. The small amount of $^1F^*_B$ that is formed persists for ~1 ns before ISC to $^3F^*_B$. This is faster than ISC in riboflavin in solution (Fig. S39d), potentially due to increased spin-orbit-coupling from the sulfur attachment on the flavin isoalloxazine ring.

**Supplementary Figure S39:** (a, b) Basis spectra of FI-4D2 (a) and riboflavin in DMSO (originally reported in Liu *et al*<sup>14</sup>) (b), extracted from the times indicated. The spectral shapes of Fl-4D2 at 20 ps and 40 ns are similar to the singlet ($^1F^*$) and triplet ($^3F^*$) spectra of riboflavin in DMSO, so are assigned $^1F_B^*$ and $^3F_B^*$. The 1 ps component of FI-4D2 is labelled $^1F_A^*$, and is attributed to an excited singlet on a different conformation that is more prone to quenching, e.g. by the protein environment or sulfur attachment. The early time contribution $^1F_A^{**}$, is attributed to vibrationally hot $^1F_A^*$ states. (c) Extracted concentrations of the spectral components in Fl-4D2, compared to the GSB decay of FI-4D2<sub>CT</sub>. For simplicity $^1F_A^*$ includes $^1F_A^{**}$ contributions. $^1F_A^*$ decays faster than the FI-4D2<sub>CT</sub> excited states, but $^1F_B^*$ lives long enough to undergo ISC. (d) Concentrations of FI-4D2 components compared to the $^1F^*$ and $^3F^*$ concentrations extracted from riboflavin in DMSO, suggesting faster ISC in the protein.

**Supplementary Figure S40:** (a) Fits of $^1F^*$ and $^3F^*$ basis spectra to the TA data of Riboflavin in DMSO at various delay times. (b) Fit of $^1F_A^*$, $^1F_B^*$, and $^3F_B^*$ to the TA data of FI-4D2 at various delay times.

###### Riboflavin in DMSO

#### FI-4D2

###### FI-4D2<sub>CT</sub>

**Supplementary Figure S41:** Jablonski diagram of photophysics observed in TA of riboflavin in DMSO solution, FI-4D2 and FI-4D2<sub>CT</sub>. NR=nonradiative and ISC = intersystem crossing.

#### S4 Organic synthesis of 7-methyl-8-chloro-10-(1'-D-ribityl)isoalloxazine

**Supplementary Figure S42:** Synthetic scheme for the synthesis of 7-methyl-8-chloro-10-(1'-D-ribityl)isoalloxazine (8-chloro-8-demethylriboflavin).

##### N-Ribityl 3-chloro-4-methyl aniline (1)

A solution of 3-chloro-4-methyl aniline (2.12 g, 15 mmol), sodium cyanoborohydride (1.89 g, 30 mmol), and D-ribose (6.75 g, 45 mmol) in methanol (30 mL) is set to reflux and stirred for 48 h. Solvent was removed under reduced pressure, and the residue was dissolved in 1 M HCl (40 mL) and swirled. A saturated solution of NaHCO<sub>3</sub> was added, and the mixture was extracted with ethyl acetate (8 × 50 mL). The combined organic layers were washed with brine and dried over MgSO<sub>4</sub>. The solvent was removed under reduced pressure to afford a brown residue. The residue was suspended in acetone, leading to precipitation of the pure product as a white solid (3.76 g, yield: 91%). <sup>1</sup>H NMR (500 MHz, d<sub>3</sub>-MeOD) δ ppm 7.01 (d, 1H, J = 8.4 Hz), 6.72 (d, 1H, J = 2.6 Hz), 6.57 (dd, 1H, J = 8.4, 2.4 Hz), 3.94 (m, 1H), 3.82 (m, 2H), 3.70 (m, 2H), 3.44 (dd, 1H, J = 12.8, 2.8 Hz), 3.15 (dd, 1H, J = 12.4, 7.6 Hz), 2.22 (s, 3H) (Fig. S43a). <sup>13</sup>C NMR (500 MHz, d<sub>3</sub>-MeOD) δ ppm 148.30, 134.14, 130.82, 123.09, 112.86, 111.84, 73.24, 72.89, 70.81, 63.13, 46.13, 17.53 (Fig. S43b). HRMS (ESI<sup>+</sup>) m/z: [M+H]<sup>+</sup> calcd. for C<sub>12</sub>H<sub>19</sub>NO<sub>4</sub>Cl 276.1003; observed 276.1001 (Fig. S44).

##### N-Ribityl-2-(4-methylphenylazo)-4-methyl-5-chloro aniline (2)

p-Toluidine (0.50 g, 4.66 mmol), 12 N HCl (1.3 mL), and H<sub>2</sub>O (3.1 mL) were combined in a 50 mL round-bottomed flask and stirred at 0 °C for 10 min. Sodium nitrite (0.33 g, 4.78 mmol) was then added slowly in small portions, and the reaction mixture was maintained at 0 °C for an additional 30 min. In a second 100 mL round-bottomed flask, N-ribityl 3-chloro-4-methylaniline 1 (1.08 g, 3.91 mmol) was dissolved in H<sub>2</sub>O (8 mL), followed by the addition of 12 N HCl (1.4 mL) and sodium acetate (1.32 g, 16.09 mmol). The mixture was cooled to -5 °C, and the solution of diazotized p-toluidine was added. The resulting mixture was stirred at -9 to -5 °C for 1 h, then at 0 °C for 2 h. After warming to 20 °C, a solution of sodium acetate (0.66 g, 8.04 mmol) in H<sub>2</sub>O (5 mL) was added slowly to maintain pH ≈ 3. The mixture was stirred at 25 °C for 16 h, affording a dark orange suspension. The solid was removed by filtration and purified by flash chromatography (DCM-MeOH 5%) to give 0.243 g (yield: 15%) of an orange solid. <sup>1</sup>H NMR (500 MHz, d<sub>3</sub>-MeOD) δ ppm 7.78 (d, 2H, J = 8.0 Hz), 7.66 (s, 1H), 7.32 (d, 2H, J = 8.5 Hz), 6.95 (s, 1H), 4.05-4.00 (m, 1H), 3.85-3.78 (m, 2H), 3.71-3.67 (m, 2H), 3.61-3.57 (m, 1H), 3.41-3.36 (m, 1H), 2.43 (s, 3H), 2.29 (s, 3H) (Fig. S45a). <sup>13</sup>C NMR (500 MHz, d<sub>3</sub>-MeOD) δ ppm 150.78, 142.98, 140.26, 138.33, 135.34, 130.26, 129.35, 122.08, 121.73, 111.76, 73.18, 73.08, 70.87, 63.28, 44.57, 20.01, 17.51 (Fig. S45b). HRMS (ESI<sup>+</sup>) m/z: [M+H]<sup>+</sup> calcd. for C<sub>19</sub>H<sub>25</sub>N<sub>3</sub>O<sub>4</sub>Cl 394.1534; observed 394.1534 (Fig. S46).

##### 7-methyl-8-chloro-10-(1'-D-ribityl)isoalloxazine (8-chloro-8-demethylriboflavin) (3)

N-Ribityl-2-(4-methylphenylazo)-4-methyl-5-chloroaniline 2 (390 mg, 0.99 mmol) with barbituric acid (201 mg, 1.57 mmol) in isobutanol (8 mL) and AcOH (2 mL) was stirred and heated at reflux for 24 h. After cooling, the solvent was removed under reduced pressure, and the residue was purified by flash chromatography (DCM-MeOH, 5%) to afford 0.135 g (yield: 30%) of a yellow solid. <sup>1</sup>H NMR (500 MHz, DMSO-d<sub>6</sub>) δ ppm 11.46 (s, 1H), 8.20 (s, 1H), 8.13 (s, 1H), 5.13 (d, 1H, J = 7.2 Hz), 4.88 (d, 1H, J = 5.3 Hz), 4.80 (d, 2H, J = 4.7 Hz), 4.68 (d, 1H,

J = 13.8 Hz), 4.47 (t, 1H, J = 5.5 Hz), 4.23 (m, 1H), 3.64 (m, 3H), 3.46 (m, 1H), 2.50 (s, 3H) (Fig. S47a). $^{13}\text{C}$ NMR (500 MHz, DMSO-$d_6$) $\delta$ ppm 160.10, 155.87, 151.28, 140.56, 138.91, 134.22, 133.49, 132.95, 118.51, 73.95, 73.18, 69.17, 63.77, 48.08, 19.61 (Fig. S47b). HRMS (ESI$^+$) m/z: $[\text{M}+\text{H}]^+$ calcd. For $\text{C}_{16}\text{H}_{17}\text{N}_4\text{O}_6\text{Cl}$ 397.0909; observed 397.0900 (Fig. S48).

**Supplementary Figure S43:** NMR characterisation of EC1 product. (a) $^1\text{H}$ and (b) $^{13}\text{C}$ NMR.

# EC1.1

BIOC-RA-42894 21 (0.372) Cm (21:31)

TOF MS ES+
7.95e8

#### Elemental Composition Report

Page 1

Tolerance = 4.0 PPM / DBE: min = -1.5, max = 80.0

Element prediction: Off

Monoisotopic Mass, Even Electron Ions

78 formula(e) evaluated with 1 results within limits (up to 50 closest results for each mass)

Elements Used:

C: 0-200 H: 0-200 N: 1-5 O: 1-5 Cl: 1-1

EC1.1

BIOC-RA-42894 21 (0.372) Cm (21:31)

TOF MS ES+
7.95e+008

**Supplementary Figure S44:** HRMS (ESI<sup>+</sup>) of the EC1 product. m/z: [M+H]<sup>+</sup> calcd. for C<sub>12</sub>H<sub>19</sub>NO<sub>4</sub>Cl 276.1003; observed 276.1001.

**Supplementary Figure S45:** NMR characterisation of EC2 product. (a) $^1\text{H}$ and (b) $^{13}\text{C}$ NMR.

# EC2.3 FC

BIOC-RA-42898 16 (0.286) Cm (13:32)

TOF MS ES+
1.26e8

#### Elemental Composition Report

Page 1

Tolerance = 4.0 PPM / DBE: min = -1.5, max = 80.0
Element prediction: Off

Monoisotopic Mass, Even Electron Ions
108 formula(e) evaluated with 1 results within limits (up to 50 closest results for each mass)
Elements Used:
C: 0-200 H: 0-200 N: 1-5 O: 1-5 Cl: 1-1
EC2.3 FC
BIOC-RA-42898 16 (0.286) Cm (13:32)

TOF MS ES+
1.26e+008

**Supplementary Figure S46:** HRMS (ESI<sup>+</sup>) of the EC2 product. m/z: [M+H]<sup>+</sup> calcd. for C<sub>19</sub>H<sub>25</sub>N<sub>3</sub>O<sub>4</sub>Cl 394.1534; observed 394.1534.

**Supplementary Figure S47:** NMR characterisation of EC3 product. (a) $^1\text{H}$ and (b) $^{13}\text{C}$ NMR.

### QE LC-Orbitrap Accurate Mass +ESI-MS

Sample: EC 3.4 FG Filename: bioc-ra-43285

bioc-ra-43285 #63-88 RT: 0.33-0.46 AV: 26 SB: 77 0.99-1.39 NL: 4.88E8
T: FTMS + p ESI Full ms [50.0000-600.0000]

| Observed m/z | Theoretical Mass | Error (ppm) | Formula | ID |
| --- | --- | --- | --- | --- |
| 397.0900 | 397.0909 | 2.27 | C<sub>16</sub>H<sub>17</sub>O<sub>6</sub>N<sub>4</sub>Cl | [M+H]<sup>+</sup> |

**Supplementary Figure S48:** HRMS (ESI<sup>+</sup>) of the EC3 product. m/z: [M+H]<sup>+</sup> calcd. For C<sub>16</sub>H<sub>17</sub>N<sub>4</sub>O<sub>6</sub>Cl 397.0909; observed 397.0900.

#### S5 Electronic Structure Benchmarking

In order to build confidence in our level of theory for the QM/MM excitation energy calculations (LR-TDDFT/TDA/ωB97X-D3/def2-svp) within our protein design pipeline, we compared results between our chosen level of theory and that of a higher level of theory, RI-CC2/def2-tzvp.<sup>15</sup> These benchmarking results were obtained for four different systems.

8-(Methylthio)lumiflavin (8-MeS-LF), a model system for the cysteinyl attached riboflavin moiety (truncated at the ribityl tail at the N10 position, and an SMe group instead of the Me group at the C8 position), was optimized on the ground electronic state to obtain $S_{0min}$ for the gas phase and a low dielectric constant to mimic a protein environment. This was performed using density functional theory (DFT) at the ωB97X-D3 functional, the def2-svp level of theory, with a polarizable continuum model (COSMO) for the mimicked protein environment in TeraChem v1.9-2023.06-dev.<sup>16–18</sup> The dielectric constant ($\epsilon$) used for the protein environment was 4.0.<sup>19</sup> 15 excited states were calculated at the same level of theory using linear-response time-dependent density functional theory employing the Tamm-Dancoff approximation (LR-TDDFT/TDA).<sup>20</sup> Resolution of identity second-order approximate coupled cluster (RI-CC2) calculations were performed for each system. These energies were calculated using Turbomole v7.8<sup>21</sup>, the COSMO implementation<sup>22</sup> for the protein environment, and a def2-tzvp basis set.

Furthermore, a sample snapshot was extracted from the Fl-4D2 QM/MM MD trajectory with the QM region being the sulfur-attached riboflavin moiety (8-dMeRF). 15 states were calculated using LR-TDDFT (LR-TDDFT/TDA/ωB97X-D3/def2-svp) and RI-CC2 (def2-tzvp) both with and without the protein environment (modelled as MM point charges). The natural transition orbitals (NTOs) were calculated using TheoDORE 2.4.2070<sup>23</sup> and visualised in ChemCraft.<sup>24</sup>

The excitation energies, oscillator strengths and transition character are displayed in tables below for all four systems. The nomenclature for the orbitals involved in the transitions is analogous to those by Marian and coworkers, see Fig.S49.<sup>25</sup> The vertical transitions obtained for 8-MeS-LF (Table S3) with LR-TDDFT/TDA/ωB97X-D3/def2-svp exhibit a larger excitation energy than the corresponding RI-CC2/def2-tzvp. The shift is more pronounced for transitions with a $\pi\pi^*$ character. Despite this rigid shift, the character of the LR-TDDFT/TDA transitions, their ordering, and the corresponding oscillator strengths align with the RI-CC2 results. Similar trends are observed upon inclusion of a dielectric constant in the calculations (Table S4), yet with a change in the character for $S_2$ and $S_3$ going from LR-TDDFT/TDA to RI-CC2 due to the destabilization of the $n_{N2} - \pi_L^*$ state and the stabilization of the $\pi_{H-1} - \pi_L^*$.

These trends are similarly observed for the 8-dMeRF moiety, without MM point charges, extracted from the Fl-4D2 QM/MM MD trajectory (Table S5). However for this set of calculations, the character of the $S_3$ and $S_4$ transitions change going from LR-TDDFT/TDA to RI-CC2, due to the stabilization of $\pi_{H-1} - \pi_L^*$ and destabilisation of the $n_{O2} - \pi_L^*$. When an environment is introduced via MM point charges (Table S6), we see the same trend as observed when moving from 8-MeS-LF in gas-phase to a dielectric constant, that is a destabilization of the $n_{N2} - \pi_L^*$ state and the stabilization of the $\pi_{H-1} - \pi_L^*$.

Overall, these results show that there is a systematic increase in the vertical excitation energies when moving from RI-CC2/def2-tzvp to LR-TDDFT/TDA/ωB97X-D3/def2-svp that is more pronounced for $\pi\pi^*$ transitions. In addition, the addition of the environment, either a dielectric constant or MM point charges, shows a consistent trend whereby the $n_{N2} - \pi_L^*$ is destabilized and $\pi_{H-1} - \pi_L^*$ is stabilized in CC2/def2-tzvp compared to LR-TDDFT/TDA/ωB97X-D3/def2-svp. Despite these systematic differences, the character of the transitions and corresponding oscillator strengths are well described in LR-TDDFT/TDA with regard to RI-CC2 and thus build confidence in our chosen level of theory to be incorporated into the design process.

$S_0 - S_1 (\pi_H - \pi_L^*)$

$S_0 - S_2 (n_{N2} - \pi_L^*)$

$S_0 - S_3 (n_{O2} - \pi_L^*)$

$S_0 - S_4 (\pi_{H-1} - \pi_L^*)$

**Supplementary Figure S49:** NTOs for low-lying transitions for 8-MeS-LF in gas-phase, calculated with RI-CC2/def2-tsvp, with orbital labels. The arrows connect the hole and particle NTOs associated with each electronic excitation.

| Electronic Transition | LR-TDDFT/TDA |  |  | RI-CC2 |  |  |
| --- | --- | --- | --- | --- | --- | --- |
| | EE (eV) | $f_{\text{osc}}$ | Character | EE (eV) | $f_{\text{osc}}$ | Character |
| S<sub>1</sub> | 3.70 | 0.607 | $\pi_H - \pi_L^*$ | 3.07 | 0.416 | $\pi_H - \pi_L^*$ |
| S<sub>2</sub> | 3.72 | 0.002 | $n_{N2} - \pi_L^*$ | 3.50 | 0.001 | $n_{N2} - \pi_L^*$ |
| S<sub>3</sub> | 4.20 | 0.000 | $n_{O2} - \pi_L^*$ | 3.70 | 0.000 | $n_{O2} - \pi_L^*$ |
| S<sub>4</sub> | 4.48 | 0.137 | $\pi_{H-1} - \pi_L^*$ | 3.77 | 0.139 | $\pi_{H-1} - \pi_L^*$ |

**Supplementary Table S3:** Excitation energies, oscillator strengths, and electronic character of the excited states for 8-MeS-LF in gas-phase for both LR-TDDFT/TDA (left) and RI-CC2 (right).

| Electronic Transition | LR-TDDFT/TDA |  |  | RI-CC2 |  |  |
| --- | --- | --- | --- | --- | --- | --- |
| | EE (eV) | $f_{\text{osc}}$ | Character | EE (eV) | $f_{\text{osc}}$ | Character |
| S<sub>1</sub> | 3.66 | 0.641 | $\pi_H - \pi_L^*$ | 2.91 | 0.569 | $\pi_H - \pi_L^*$ |
| S<sub>2</sub> | 3.85 | 0.002 | $n_{N2} - \pi_L^*$ | 3.51 | 0.159 | $\pi_{H-1} - \pi_L^*$ |
| S<sub>3</sub> | 4.30 | 0.139 | $\pi_{H-1} - \pi_L^*$ | 3.61 | 0.001 | $n_{N2} - \pi_L^*$ |
| S<sub>4</sub> | 4.41 | 0.000 | $n_{O2} - \pi_L^*$ | 3.91 | 0.000 | $n_{O2} - \pi_L^*$ |

**Supplementary Table S4:** Excitation energies, oscillator strengths, and electronic character of the excited states for 8-MeS-LF in a dielectric constant where $\epsilon = 4$ for both LR-TDDFT/TDA (left) and RI-CC2 (right). The characters of the transitions are the same as in the gas phase and so the nomenclature is maintained.

| Electronic Transition | LR-TDDFT/TDA |  |  | RI-CC2 |  |  |
| --- | --- | --- | --- | --- | --- | --- |
| | EE (eV) | $f_{\text{osc}}$ | Character | EE (eV) | $f_{\text{osc}}$ | Character |
| S<sub>1</sub> | 3.42 | 0.424 | $\pi_H - \pi_L^*$ | 2.73 | 0.274 | $\pi_H - \pi_L^*$ |
| S<sub>2</sub> | 3.53 | 0.072 | $n_{N2} - \pi_L^*$ | 3.15 | 0.001 | $n_{N2} - \pi_L^*$ |
| S<sub>3</sub> | 3.87 | 0.011 | $n_{O2} - \pi_L^*$ | 3.28 | 0.202 | $\pi_{H-1} - \pi_L^*$ |
| S<sub>4</sub> | 4.07 | 0.193 | $\pi_{H-1} - \pi_L^*$ | 3.37 | 0.017 | $n_{O2} - \pi_L^*$ |

**Supplementary Table S5:** Excitation energies, oscillator strengths, and electronic character of the excited states for a snapshot from the Fl-4D2 QM/MM MD trajectory for both LR-TDDFT/TDA (left) and RI-CC2 (right). These calculation were performed without the MM point charges. The orbitals are not identical to those from the calculations performed on 8-MeS-LF, due to presence of the ribityl tail, however the nomenclature is maintained as the orbitals involved in the transitions are analogous if not identical.

| Electronic Transition | LR-TDDFT/TDA |  |  | RI-CC2 |  |  |
| --- | --- | --- | --- | --- | --- | --- |
| | EE (eV) | $f_{\text{osc}}$ | Character | EE (eV) | $f_{\text{osc}}$ | Character |
| S<sub>1</sub> | 3.40 | 0.459 | $\pi_H - \pi_L^*$ | 2.71 | 0.317 | $\pi_H - \pi_L^*$ |
| S<sub>2</sub> | 3.53 | 0.072 | $n_{N2} - \pi_L^*$ | 3.13 | 0.125 | $\pi_{H-1} - \pi_L^*$ |
| S<sub>3</sub> | 3.95 | 0.191 | $\pi_{H-1} - \pi_L^*$ | 3.16 | 0.066 | $n_{N2} - \pi_L^*$ |
| S<sub>4</sub> | 4.28 | 0.001 | $n_{O2} - \pi_L^*$ | 3.76 | 0.001 | $n_{O2} - \pi_L^*$ |

**Supplementary Table S6:** Excitation energies, oscillator strengths, and electronic character of the excited states for a snapshot from the Fl-4D2 QM/MM MD trajectory, with MM point charges, for both LR-TDDFT/TDA (left) and RI-CC2 (right). The orbitals are not identical to those from the calculations performed on 8-MeS-LF, due to presence of the ribityl tail, however the nomenclature is maintained as the orbitals involved in the transitions are analogous if not identical.

#### S6 Supplementary Methods

##### S6.1 Mass Spectrometry

Electrospray ionisation (ESI) MS was performed on Waters Xevo G2-XS QTOF LC-MS, without a column for liquid chromatography. Protein samples of 50-100 $\mu\text{M}$ were exchanged into 100 mM ammonium acetate and injected at a flow rate of 0.25 mL/min; $m/z$ values were obtained in positive ion mode, with the capillary and cone voltages set to 3.6 kV and 50 V respectively. Spectral deconvolution was achieved using MaxEnt1 software.<sup>26</sup>

##### S6.2 Circular Dichroism (CD) Spectroscopy

CD spectra were obtained using a JASCO J-1500 spectrophotometer, with protein concentrations of 5-12 $\mu\text{M}$ and a prior blank recorded with buffer. Raw ellipticity (mdeg) was recorded at 222 nm at 1°C intervals from 5-95 °C, with eight repeats of full spectra (190-250 nm) obtained at the following temperatures (°C): 5, 25, 35, 55, 75 and 95. Refolding studies were performed at the same intervals with decreasing temperatures. The average signal was converted to mean residue ellipticity (MRE) using Eq. S12, whereby $n$ is the number of peptide bonds present in the protein, $l$ is the pathlength and $c$ is the protein concentration.

$$\text{MRE}(\text{deg} \cdot \text{cm}^2 \cdot \text{dmol}^{-1}) = \frac{\theta(\text{mdeg}) \cdot 10^6}{l(\text{mm}) \cdot c(\mu\text{M}) \cdot n} \quad (\text{S12})$$

##### S6.3 Steady-state Absorption and Fluorescence Spectroscopy

UV/Visible absorption spectra were recorded with a Cary 60 UV/Visible Spectrophotometer (Agilent Technologies) with a scan rate of 2000 nm/min and a prior baseline with the corresponding sample buffer. Fluorescence emission spectra were recorded with a Cary Eclipse Fluorescence Spectrophotometer (Agilent Technologies), with an excitation wavelength of 460 nm and 10 nm excitation/emission slit widths.

##### S6.4 Optically Transparent Thin-Layer Electrochemistry (OTTLE)

OTTLE was used to spectroscopically determine redox potentials of the bound cofactors upon potentiometric titration. Inside a modified EPR cuvette of 0.1 cm pathlength, protein sample was made up to an OD at $\lambda_{\text{max}}$ of approximately 0.5-0.7 alongside 10% v/v glycerol and the following redox mediators: 20 $\mu\text{M}$ phenazine ethosulfate, 50 $\mu\text{M}$ duroquinone, 6 $\mu\text{M}$ indigotrisulfonate, 25 $\mu\text{M}$ 2-hydroxy-1,4-naphthoquinone, 20 $\mu\text{M}$ phenazine and 20 $\mu\text{M}$ anthroquinone-2-sulfonate. Ag/AgCl reference (RE-5B, BASi), platinum counter, and platinum working electrodes were added to the cell and the remaining chamber filled with buffer. Potential was applied to the cell through a BioLogic SP-150 potentiostat and spectra recorded after 30 min equilibration using a Cary 60 UV/Visible spectrophotometer.

Absorbance at wavelengths associated with cofactor redox states (429 nm for oxidized heme B Soret and 458, 485, and 462 nm for oxidized flavin in Fl-4D2, FC-4D2 and FI-4D2<sub>CT</sub> respectively) were plotted against applied potential, shifted with a calibration of ~225 mV to be against Standard Hydrogen Electrode (SHE). The sigmoidal traces were then fit to derivatives of the Nernst equation to determine the midpoint potential; heme titrations were fit to a $1e^-$ equation (Eq. S13), whilst the flavin traces were fit with either a sequential $2 \times 1e^-$ (Eq. S14) or a concerted $2e^-$ equation (Eq. S15).

$$\text{Abs} = \frac{A + \left( B \cdot 10^{\frac{Em-V}{0.059}} \right)}{1 + 10^{\frac{Em-V}{0.059}}} \quad (\text{S13})$$

$$\text{Abs} = \frac{\left( A \cdot 10^{\frac{V-E1}{0.059}} \right) + B + \left( C \cdot 10^{\frac{E2-V}{0.059}} \right)}{1 + 10^{\frac{V-E1}{0.059}} + 10^{\frac{E2-V}{0.059}}} \quad (\text{S14})$$

$$\text{Abs} = \frac{A + \left( B \cdot 10^{\frac{Em-V}{0.0295}} \right)}{1 + 10^{\frac{Em-V}{0.0295}}} \quad (\text{S15})$$

#### S6.5 Rapid Freeze Quench EPR Spectroscopy

Rapid freeze-quenching (RFQ) of the EPR samples was performed on an isopentane-free apparatus as described before.<sup>27</sup> An Update Instrument (Madison, WI) mixing machine and a home-built apparatus were used for freezing the ejected mixtures on the surface of a rapidly rotating aluminum disk kept at liquid nitrogen temperature. 50 μM FI-4D2 and 5 mM sodium dithionite solutions were kept at room temperature in the Update Instrument freeze-quenching apparatus and were mixed in the mixing chamber. A 100 ms timepoint sample was made using a calibrated aging hose. The sample was frozen on the cold disk and transferred to a funnel attached to an EPR tube and packed with a Teflon-tipped stainless steel rod.

EPR spectra were recorded on a Bruker EMX (X-band) EPR spectrometer with the use of an Oxford Instruments liquid-helium cooling system and a spherical high-quality ER 4122 (SP 9703) Bruker resonator. EPR spectra were measured with a temperature of 120 K, microwave frequency of 9.36 GHz, microwave power of 1 μW, modulation frequency of 100 kHz, modulation amplitude of 5 G, time constant of 80 ms, and 20 scans. EPR spectra were processed using Kazan Viewer by Alexey Silakov (Pennsylvania State University) and EPR spectral simulations were performed using EasySpin<sup>28</sup> in the MATLAB environment.

#### S6.6 Protein Nuclear Magnetic Resonance (NMR)

<sup>15</sup>N isotopically-labelled recombinant protein was expressed following the protocol listed by Marley *et al*<sup>29</sup> before purification as listed above. <sup>1</sup>H-<sup>15</sup>N SOFAST-HMQC spectra were acquired on a uniformly <sup>15</sup>N-labelled protein using the SOFAST-HMQC pulse sequence developed by Schanda and Brutscher.<sup>30</sup> Spectra were recorded at 298K on a Bruker Avance III HD spectrometer operating at 700MHz with a 1.7 mm TCI microcryoprobe using the standard Bruker pulse program before being processed with Topspin 3.6.1.
